# Fables and foibles: a critical analysis of the Palaeoflora database and the Coexistence approach for palaeoclimate reconstruction

**DOI:** 10.1101/016378

**Authors:** Guido W. Grimm, Johannes M. Bouchal, Thomas Denk, Alastair J. Potts

**Affiliations:** University of Vienna, Department of Palaeontology, Wien; Swedish Museum of Natural History, Department of Palaeobiology, Stockholm; Nelson-Mandela Metropolitan University, Botany Department, Port Elizabeth

**Keywords:** coexistence approach, mutual climate range techniques, nearest-living relative concept, Palaeoflora database, data error

## Abstract

The “Coexistence Approach” is a mutual climate range (MCR) technique combined with the nearest-living relative (NLR) concept. It has been widely used for palaeoclimate reconstructions based on Eurasian plant fossil assemblages, most of them palynofloras (studied using light microscopy). The results have been surprisingly uniform, typically converging to subtropical, per-humid or monsoonal conditions. Studies based on the coexistence approach have had a marked impact in literature, generating over 10,000 citations thus far. However, recent studies have pointed out inherent theoretical and practical problems entangled in the application of this widely used method. But so far little is known how results generated by the coexistence approach are affected by subjective errors, data errors, and violations of the basic assumptions. The majority of Coexistence Approach studies make use of the Palaeoflora database (the combination of which will be abbreviated to CA+PF). Testing results produced by CA+PF studies has been hindered by the general unavailability of the contents in the underlying Palaeoflora database; two exceptions are the mean-annual temperature tolerances and lists of assigned associations between fossils and nearest-living relatives. Using a recently published study on the Eocene of China, which provides the first usable insight into the data structure of the Palaeoflora database, we compare the theory and practice of Coexistence Approach using the Palaeoflora database (CA+PF). We show that CA+PF is riddled by association and climate data error. We reveal the flaws in the application of CA, which is often in stark contrast to the theory of the method. We show that CA+PF is highly vulnerable against numerous sources of errors, mainly because it lacks safeguards that could identify unreliable data. We demonstrate that the CA+PF produces coherent, pseudo-precise results even for artificially generated, random plant assemblages. Alternative MCR-NLR methods can surpass the most imminent deficits of CA, and may be used as a stop-gap until more accurate bioclimatic and distribution data on potential Eurasian NLRs, and theoretically- and statistically-robust methods will become available. Finally, general guidelines are provided for the future application of methods using the mutual climatic range with nearest living relatives approach when reconstructing climate from plant fossil assemblages.

## 1 Introduction

Reconstructing past climates is an extensive field of research where a range of proxies are used to determine this aspect of the earth’s history. There are many proxies used to determine past climates, ranging from stable isotopes from ocean floor sediments (e.g. Zachos et al., 2001) to biological proxies such as pollen (e.g. Zagwijn, 1994). The approaches used to reconstruct past climates need to be theoretically and methodologically sound else results may be erroneous and misleading. The “Coexistence Approach” has been widely used by to reconstruct Cenozoic climates from Eurasian plant fossils (Utescher et al., 2014), generating in total over 10,000 citations. Here we examine whether the Coexistence Approach is scientifically sound in terms of its methodological approach; in a separate study we consider the theoretical underpinnings of the Coexistence Approach and other methods based on the mutual climate range coupled with nearest living relative associations. In order to provide this assessment we will focus on two recent Coexistence Approach publications: the first is a review of the method by Utescher et al. (2014), which also provides new guidelines for best practise, and the second is a study that reconstructs the Eocene climate of China (Quan et al., 2012) using the Palaeoflora database (Coexistence Approach studies that use the Palaeoflora database are henceforth abbreviated to CA+PF); the authors consider the latter to be the most extensively documented CA+PF study to date. The data (climatic ranges and assignments of nearest living relatives) for previous studies using this method have, by and large, been unavailable for scrutiny (Grimm and Denk, 2012). We argue that if we find that the method and application of the Coexistence Approach is severely flawed based on an assessment of these two publications, then climate reconstructions generated by all other such studies over the last 15 years should be considered spurious. We also provide a brief summary of alternative approaches and suggestions for the future of this field.

The Coexistence Approach is a mutual climate range technique in the family of “nearest-living relative” approaches. Using nearest-living relatives is usually obligatory as most of the species present during the Cenozoic are now extinct. The foundational principle behind methods that use the mutual climate range and nearest-living relative approaches (henceforth referred to as MCR-NLR) is that of physiological uniformitarianism (Tiffney, 2008): this is the assumption that form does not change as long as a lineage thrives in the same ecological or climatic niche. Mutual climate range techniques share the basic idea of species distribution modelling: the climatic niche or tolerance of a species (sensu Hutchinson, 1957) can be extracted from its current distribution and is stable over time. Species distribution modelling estimates an n-dimensional niche for a single species, whereas mutual climate range techniques estimate the mutually shared climatic range of a community, however the niche dimensions are usually analysed in isolation or with limited interactions. Pure mutual climate range techniques are restricted to the relative recent past, where an assemblage consists of extant taxa (e.g. Böcher, 1995; Elias, 2001; Thompson et al., 2012a). The nearest-living relative principle extends such approaches to fossil taxa; this principle has been used since the dawn of palaeobotanical research (Taylor et al., 2009, p. 6). It involves replacing each fossil species by a modern analogue representing the same phylogenetic lineage. The basic assumption of MCR-NLR is that the climatic niche has not substantially changed for *a phylogenetic lineage*, including the fossil and its nearest-living relative.

In case of the Coexistence Approach, the MCR-NLR estimates are based on minimum-maximum tolerances recorded for modern taxa, the nearest-living relatives, which then are used to reconstruct the putative mutual climate ranges for the fossil (usually plant) assemblage (Mosbrugger and Utescher, 1997; Utescher et al., 2014). Each individual climate parameter is estimated independently. Aside from theoretical issues, which will be addressed elsewhere, this has many potential sources of subjective and data errors (Mosbrugger and Utescher, 1997; Grimm and Denk, 2012; Utescher et al., 2014). The most apparent are misidentification of fossil specimens, poor choice of nearest-living relatives and difficulties in obtaining accurate tolerance data for these taxa. The applicability of the method and the validity of the Palaeoflora database (Utescher and Mosbrugger, 2009), which has been used in nearly all Coexistence Approach studies, has rarely been questioned (but see Klotz, 1999; Grimm and Denk, 2012). However, Utescher et al. (2014) have recently reassessed the Coexistence Approach and provided extensive details on potential errors. The majority of these have thus far been largely ignored in the Coexistence Approach literature (but see Eldrett et al., 2014; Kotthoff et al., 2014). Earlier, Grimm and Denk (2012) pointed out similar inconsistencies and shortcomings of this method for the reconstruction of mean-annual temperature using the Palaeoflora database and provided a number of suggestions on how to correct for these. The conclusions and suggestions put forward by Utescher et al. (2014) largely overlap with this earlier study (**Table 1**), suggesting that these authors independently followed the same lines of reasoning after 15 years of uncritically using this method on Eurasian plant fossil assemblages.

**Table 1.** Comparison of some points raised by Grimm and Denk (2012), the standard practise in Coexistence Approach studies using the Palaeoflora databse (CA+PF) from 1997 until 2013, and Utescher et al.’s (2014) recommendations.

| Issue | Grimm and Denk (2012) | CA+PF literature prior to 2012 | Utescher et al. (2014) |
| --- | --- | --- | --- |
| Documentation | Full documentation and disclosure is necessary | In most cases just climate reconstructions, sometimes with lists of fossil taxa and/or nearest-living relatives. No tolerance data<sup>a</sup> | Fossil lists, nearest-living relative associations and tolerance data should be documented |
| Misidentified fossils | Are a problem; examples provided | Are not discussed, critical fossils rarely figured | Give rise to unavoidable “uncertainties” |
| Wrong or unrepresentative NLRs | Are a problem; examples provided | Are not discussed, but frequently included | Give rise to unavoidable “uncertainties” |
| Maximum possible accuracy of nearest-living relative (extant taxa) climate tolerances | 1–2 °C for temperature data | No not discussed | Maximum of 1 °C for temperature data, and 100 mm/year for precipitation<sup>b</sup> |
| Gridded climate data, in areas with few stations and/or strong topographical relief. | Can be problematic; examples provided and means of corrections (e.g. verifying application of the adiabatic lapse rate in a region of interest using station data) | Not used | Can be problematic, following the same logic outlined by Grimm and Denk (2012) |
| Gridded-climate, multi-station data vs. hand-picked 4–6 ‘extreme’ stations (basis for PF NLR tolerances) | Gridded-climate (corrected for altitudinal effects) and multi-station data outperform PF tolerances, as seen in CA intervals reconstructed for floras growing in penultimate vicinity of climate stations | – | Careful selection of 4–6 ‘extreme’ stations can compensate for deficits of gridded-climate data (no evidence produced); altitudinal information should be considered, essentially in the way Grimm and Denk (2012) did it |
| Maximum precision (resolution) of the Coexistence Approach | Rarely 3 °C, typically $\geq 5$ °C for MAT using realistic tolerances; defined based on more than 200 *modern validation floras* | Studies report climate intervals with (pseudo-) precision down to 0 °C, <10 mm/year or month and discuss shifts in climate of $\leq 2$ °C and < 100 mm/year or 10 mm/month | 2.1 ($\pm 1.35$) °C for MAT; defined based on *CA/PF results* of 357 Cenozoic floras (29 taxa in mean); CA/PF results should not be interpreted at the precision that is reported<sup>c</sup> |
| Accuracy of tolerance data in the Palaeoflora database | Fairly to highly inaccurate. For example, two thirds of ~350 nearest-living relatives (species, genera, families) tolerated MAT that were at least 2° lower than recorded in the Palaeoflora database; one third had lower MAT tolerances that were more than 5 °C too warm (using conservative estimates) | Considered highly or sufficiently accurate following the validation by Mosbrugger and Utescher (1997)<sup>d</sup> | Recorded MAT ranges cover 65.05–103.39% of the MAT ranges reported by Thompson et al. (1999b) for 25 North American species, with little/no relevance for application of CA |

One of the primary criticisms raised by Grimm and Denk (2012) related to the poor documentation in the majority of CA+PF studies (**Table 1**). The lack of documentation for climate tolerances of the nearest-living relatives is particularly problematic as Utescher et al. (2014) state that the Palaeoflora database undergoes permanent updates; thus comparing published results is impossible. Utescher et al. (2014) now recommend that in the future – after 15 years of “successfully” applying this method – all used data (lists of fossils and nearest-living relative associations, tolerance data of nearest-living relatives) shall be documented. The only pre-2014 publications that are in-line with the new guidelines are those that used (to some degree) independent climate tolerance data (i.e. not extracted from the Palaeoflora database; Xu et al., 2008; Jacques et al., 2011).

Mosbrugger and Utescher (1997) list four basic assumptions that need to be fulfilled to apply the Coexistence Approach (**Table 2**; cf. Utescher et al., 2014, section 3): 1) a nearest-living relative can be identified, 2) the minimum and maximum climate tolerances of the fossil taxon are similar to that of its nearest-living relative, 3) these tolerances of the nearest-living relative can be deduced from its area of distribution, and 4) the modern climate [and distribution] data are of good quality. Because of poor documentation, it has been impossible to assess whether any violations of these basic assumptions bias the outcome of CA+PF results. The only parameter that has been evaluated is the mean annual temperature, which showed little reliability of palaeoclimate reconstructions based on the Coexistence Approach (Grimm and Denk, 2012). In 2012, a CA+PF study was published that complies (almost entirely) with the new policy (Quan et al., 2012). Using literature data, Quan et al. compiled taxon lists for the Eocene of China, the distribution of these assemblages span the entire country. Please note that we use the climate parameter abbreviations common to Coexistence Approach studies and these are listed in **Section 1.1**. Based on the substantial drop observed in CMT (average temperature of the coldest month), slight potential increase in WMT (average temperature of the warmest month) and generally low LMP (monthly precipitation of the driest month) observed in southern China, they concluded that the Himalayan uplift must have already occurred by the late Eocene inflicting monsoonal climate in southern China. Appendix A of Quan et al. (2012) lists the floral assemblages, their constituent fossil taxa and the nearest-living relatives used to represent those taxa. Appendix B of Quan et al. (2012) lists the climatic min-max tolerances of all nearest-living relatives (including unused taxa) extracted from Palaeoflora database. The number of nearest-living relatives per assemblage is high (usually more than 20 up, with one instance over 100). Therefore, one would expect that the reconstructed intervals are highly reliable (Utescher et al., 2014, p. 61). The study of Quan et al. (2012) provides the first opportunity to compare theory (Mosbrugger and Utescher, 1997; Utescher et al., 2014) and practise of Coexistence Approach studies relying on the Palaeoflora database. Utescher et al. (2014, p. 60) state that the “*general agreement*” of the results from this method with alternative palaeoclimate reconstructions (using floristic assemblages or isotope data) and palaeoclimate model predictions demonstrates that (inevitable or correctable) errors (“uncertainties”, Utescher et al., 2014) in the database are not of general concern. Here we will examine this claim using the data of Quan et al. (2012).

**Table 2.** Assumptions, guidelines and terminology of the Coexistence Approach (CA)

|  |
| --- |
| *Assumptions* |
| 1) A nearest-living relative (NLR) can be identified. |
| 2) The minimum and maximum climate tolerances of the fossil taxon are similar to that of its NLR. |
| 3) The climate tolerances of the NLR can be deduced from its area of distribution. |
| 4) The modern climate [and distribution] data are of good quality. |
| *Guidelines for Coexistence Approach* |
| A) Taxa with a much larger distribution in the past should be excluded |
| B) Monotypic genera should be excluded |
| C) Cosmopolitan and aquatic taxa should be excluded |
| D) If not all NLR have overlapping tolerances, i.e. a mutual climate range, the coexistence interval is defined by the mutual climate range of the highest number of potentially coexisting NLRs. This may lead to ‘ambiguous intervals’, indicative for a mixed flora (according Utescher et al., 2014) or recognition of one or few NLRs as ‘climatic outliers’, indicative of a violation of at least one of the four basic assumptions (according Mosbrugger and Utescher, 1997; Utescher et al., 2014). |
| *Key terms* |
| **Mutual climate range (MCR)** – the mutually shared, 1- or 2-dimensional climate space by two or more organisms |
| **Nearest-living relatives (NLR)** – The extant, phylogenetically controlled, analogue of a fossil taxon. In a strict sense, a NLR is an extant species that can be considered to be a direct relative (descendant) of a fossil species. In practise, most NLRs are systematically defined, i.e. an extant genus is taken as NLR for a fossil species with systematic-phylogenetic affinities to this extant genus. |
| **Coexistence Approach (CA) interval (Fig. 1A)** – the mutual climate range of the maximum-possible number of NLRs (not necessarily all). |
| **Ambiguous CA interval (Fig. 1B)** – Two or more alternative CA intervals referring to the mutual climate ranges of equally large subsets of NLRs, in cases where not all NLR have a mutually shared climate range |
| **Climatic outlier (Fig. 1C)** – A NLR that is excluded from the calculation of CA intervals following the procedure outlined in Mosbrugger and Utescher (1997) to define CA intervals in cases where not all NLR have a mutually shared climate range |

**Figure 1.**
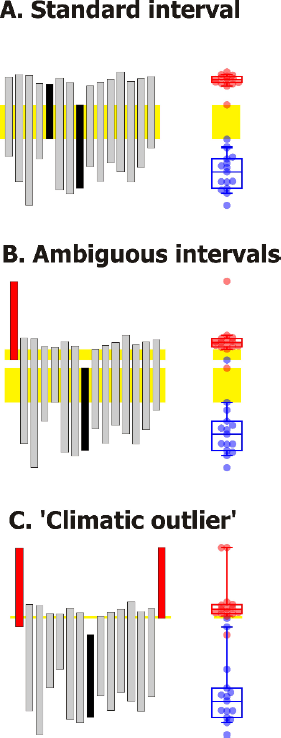
Definition of intervals and ‘climatic outliers’ according to the ‘Coexistence Approach’. **A.** Assemblage with no apparent outliers; the coexistence interval (light green) for the assemblage is defined by the mutually shared climate range of the two least tolerant (with the lowest maximum, highest minimum tolerance, respectively) nearest-living relatives (black). **B.** The assemblage differs from **A** by the occurrence of an exotic nearest-living relative (red bar), which is a statistical outlier. Two ‘ambiguous’ coexistence intervals (light green) are the result, each excluding one taxon (black or red) as an outlier. **C**. Two nearest-living relatives (red bars) trigger the recognition of the black nearest-living relative as ‘climatic outlier’. Note that the nearest-living relatives highlighted in red are likely exotic members of the assemblage as they share a small proportion of the climate space with the rest. Nonetheless, as highlighted in C, a few exotic nearest-living relatives can overrule alternative, likely more likely, climate reconstruction scenarios.

### 1.1 Abbreviations

The following abbreviations will be used throughout the text:

CA+PF = coexistence approach using the Palaeoflora database MCR = mutual climate range

MAT = mean annual temperature

CMT = mean temperature of the coldest month

WMT = mean temperature of the warmest month

MAP = mean annual precipitation [mm/year]

HMP = monthly precipitation of the wettest month

LMP = monthly precipitation of the driest month

WMP = monthly precipitation of the warmest month

## 2 Identification of nearest living relatives

### 2.1 The theory

Ideally, only one or several modern species would be considered the closest (nearest-living) relative\s of one fossil taxon. However, many species-level associations are problematic (Grimm and Denk, 2012) and studies that clarify systematic affinities of fossil taxa are either not available or generally overlooked by researchers using the Coexistence Approach and/or the curators of the Palaeoflora database (Grimm and Denk, 2012). Examples of overlooked updated taxa critical to palaeoclimate reconstructions include Engelhardioideae (Manchester, 1987; Manos et al., 2007) and the Fagaceae genera *Fagus* and *Quercus* (Denk et al., 2002, 2005; Grimm et al., 2007; Denk and Grimm, 2009a, b, 2010). According to Utescher et al. (2014), the Palaeoflora database includes association information of c. 5800 macrofossil taxa and c. 2500 microfossil taxa (ranging across species, genera and families), but this information cannot be tested based on available documentation. The homepage lists numerous palaeobotanical works which have been harvested for potential associations between fossil taxa and nearest-living relatives, but this information is not linked to the association list. This linkage is essential as Grimm and Denk (2012) and Utescher et al. (2014, p. 64) warn against the blind usage of such association lists, in particular when using macrofossil assemblages. Indeed, Grimm and Denk (2012) highlighted erroneous associations that severely biased Coexistence Approach reconstructions. Utescher et al. (2014) stated that good taxonomy is critical for CA+PF studies, and that efforts were underway to incorporate new phylogenetic and systematic-taxonomic evidence into the Palaeoflora database. Often it is even difficult to determine in CA+PF studies to what degree the fossil taxon–nearest-living relative association follows the Palaeoflora database or is based on other work, which adds additional uncertainty.

### 2.2 The reality

In most cases, Quan et al. (2012) selected a genus-level nearest-living relative based on the (assumed) generic affinity of the fossil taxon (macrofossil or microfossil). If the generic affinity of a fossil taxon was ambiguous, Quan et al. (2012) used the subfamily or family. If the fossil taxon resembled more than one (related or unrelated) modern genera or families then an artificially fused taxon was created by merging the tolerances of the modern taxa. This violates assumption 1 of the Coexistence Approach (**Table 2**): a nearest living relative can be identified (outlined below; see also Grimm and Denk, 2012).

Poor taxonomic control can be a source of error, particular in studies that use macrofossil assemblages to reconstruct past climates. Utescher et al. (2014, p. 64) mystically claim that the “*type status*” of pollen “*accommodates [taxonomic] uncertainties*”. Most of the assemblages used by Quan et al. (2012) are palynofloras studied using light microscopy – not scanning-electron microscopy that could provide better taxonomic resolution – translated into lists of genus- to family-level nearest-living relatives. Unfortunately, the translation of fossil pollen taxa does *not include the original ‘species’ names*. This is crucial for the choice of a particular nearest-living relative as palynological taxonomy focuses on *form* and not *systematic affinity*. Thus, ‘species’ of the same pollen ‘genus’ may represent different families or higher-level taxa. For example, Quan et al. (2012) link *Inaperturopollenites* with “taxodioid Cupressaceae” (former Taxodiaceae). However *Inaperturopollenites* ‘species’ can represent a far wider range of biological taxa (**Table 3**). We observe that, except for very few examples, pollen genera were linked with a specific nearest-living relative by Quan et al. (2012). Consequently, we are forced to assume that the omitted species names are irrelevant in case of the Chinese Eocene floras. Under this assumption and using available compendia on palynological taxa (Martin and Drew, 1969; Huang, 1972; Solomon et al., 1973; Stuchlik et al., 2001; Stuchlik et al., 2002; Beug, 2004; Stuchlik et al., 2009; Li, 2011; Miyoshi et al., 2011; Stuchlik et al., 2014) we crosschecked the validity of the associations used to link fossil taxa with a nearest-living relative.

**Table 3.** Botanical affinities of ‘species’ of the pollen ‘genus’ *Inaperturopollenites.* Species that have affinities to “Taxodiaceae” (paraphyletic group including Sequoioideae and Taxodioideae) in bold.

| Taxon | Author, year | Affinity | Reference<sup>a</sup> |
| --- | --- | --- | --- |
| *Inaperturopollenites ahikokuensis* | Takahashi 1962 | Salicaceae (*Populus*),<br>Triuridaceae (*Sciaphila*) | CFS&P, 25-120 |
| ***I. concedipites*** | (Wodehouse, 1933)<br>Krutsch 1971 | "Taxodiaceae"<br>(*Taxodium*,<br>*Glyptostrobus*) | Stuchlik et al., 2002, p.<br>50, Plate 71, figs 9-29 |
| ***I. crassatus*** | Takahashi 1961 | "Taxodiaceae" | CFS&P, 21-063 |
| ***I. dubius*** | (Potonié & Venitz,<br>1934) Thomson &<br>Pflug 1953 | "Taxodiaceae" | Stuchlik et al., 2002, p.<br>51, Plate 71, figs 1-8 |
| *I. falsus* | Takahashi 1964 | ? | CFS&P, 36-187 |
| *I. giganteus* | Góczán 1964 | ? | CFS&P, 33-083 |
| *I. globulus* | Weyland & Greifeld<br>1953 | ? | CFS&P, 02-056 |
| *I. globulus* | Nilsson 1958 | Pseudo-*Corylus* | CFS&P, 24-038 |
| *I. heskemensis* | Krivánné-Hutter 1961 | ? | CFS&P, 21-024 |
| *I. immutatus* | Takahashi 1961 | Aristolochiaceae | CFS&P, 21-064 |
| *I. insignis* | Manum 1962 | *Larix*, *Pseudotsuga* | CFS&P, 25-044 |
| *I. irregularis* | Nilsson 1958 | ? | CFS&P, 24-039 |
| ***I. ligularis*** | Takahashi 1961 | *Sequoia* or *Metasequoia* | CFS&P, 21-065 |
| *I. limbatus* | Balme 1957 | ? | CFS&P, 16-141 |
| *I. orbiculatus* | Nilsson 1958 | ? | CFS&P, 24-040 |
| *I. parviundulatus* | Takahashi 1964 | ? | CFS&P, 36-188 |
| *I. parvoglobulus* | Weyland & Greifeld<br>1953 | ? | CFS&P, 02-057 |
| *I. parvus* | Takahashi 1963 | ? | CFS&P, 25-133 |
| *I. patellaeformis* | Weyland & Greifeld<br>1953 | ? | CFS&P, 02-055 |
| *I. pseudosetarius* | Weyland & Pflug 1957 | *Smilax*, *Hydrocharis* | CFS&P, 25-139 |
| *I. redi* | de Jersey 1959 | ? | CFS&P, 26-036 |
| *I. sublevis* | Nilsson 1958 | ? | CFS&P, 24-041 |
| *I. triangularis* | Nilsson 1958 | ? | CFS&P, 24-042 |
| *I. undulatus* | Weyland & Greifeld<br>1953 | ? | CFS&P, 02-059 |
| *I. verrucosus* | Weyland & Greifeld<br>1953 | ? | CFS&P, 02-060 |
| ***I. verrupapillatus*** | Trevisan 1967 | "Taxodiaceae"<br>(*Taxodium*,<br>*Glyptostrobus*) | Stuchlik et al., 2002, 52,<br>Plate72, figs 6-11;<br>Plate73, figs 1-8 |

When screening the CA+PF documentation provided by Quan et al. (2012) the following can be observed: 14 nearest-living relatives chosen by the authors are invalid because they are *(i)* combinations of unrelated taxa (discussed below), *(ii)* refer to obsolete, polyphyletic taxa, or *(iii)* have been linked to a wrong nearest-living relative (violation of assumption 1, **Table 2**). For examples, “*Engelhardtia*”, “*Engelhardtioidites*”, and “*Engelhardtioipollenites*” have been erroneously linked with extant *Engelhardia* and “*Cupuliferoipollenites*” with extant “Castaneae” (Box 1). Twenty nearest-living relatives are highly problematic because the chosen extant genus or family is not the only one with such a type of pollen; exact systematic affinity cannot be unambiguously identified using a light microscope and, in some cases, not even using scanning electron microscopy. But does this affect the Coexistence Approach reconstructions? In **Table 4** we give examples of erroneous or problematic nearest-living relative associations that directly influenced the reconstructed climate intervals reported by Quan et al. (2012); **Figure 2** illustrates this at the example of the infamous *“Engelhard(t)ia*”.

**Figure 2.**
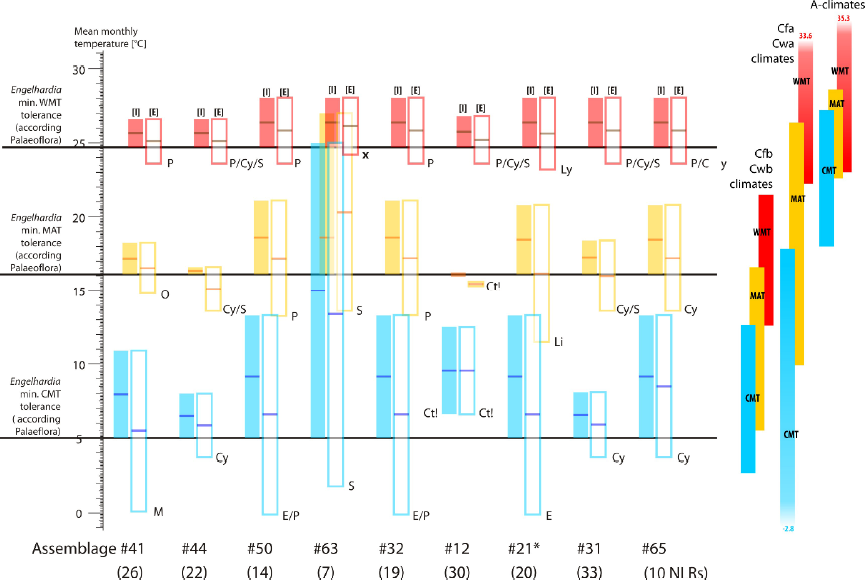
Examples of reconstructed climate intervals reported in Quan et al. (2012) that were affected by the invalid (see text) nearest-living relatives, *Engelhardia*: [I]ncluded versus [E]xcluded from the reconstruction. The temperature ranges for major climate zones in East Asia and tropical Australasia are shown for reference. The removal of *Engelhardia* often resulted in the climate interval being determined by another problematic nearest-living relative; these are highlighted (subscripts): Ct! = Cyatheaceae (highly diverse tropical to southern temperate fern family, with MATmin and WMTmin at least 5 °C too high, CMTmin at least 1.5 °C too high; **File S2** in online supporting archive); Cy = Cyrillaceae (tropical Ericales family endemic to the Caribbean); E = *Ephedra* (+Ephedraceae, monotypic family, recorded with different MATmin and WMTmin); Li = *Liquidambar* (genus with three highly disjunct, relict species); Ly = Lygodiaceae (monotypic family, recorded WMTmin > recorded WMTmin of its constituent genus); M = Myrtaceae; O = Olacaceae (pantropical family, extending marginally into subtropics); P = *Planera* (monotypic Ulmaceae genus of North America, geographically restricted, problematic fossil–nearest-living relative association), S = Symplocaceae (monotypic Ericales family; recorded tolerances in strong conflict with distribution of modern species in northeast Asia), **x** = flawed, artificial nearest-living relative (**Table 5**).

**Table 4.** List of invalid **[I]** and problematic **[P]** nearest-living relatives used in the study of Quan et al. (2012) that either define coexistence intervals or are recognised as “climatic outliers” for a given climate parameter. The influence of the nearest living relative on the interval is specified as: no affect (-), defines the lower boundary (LB) or the upper boundary (UB), or was labelled an outlier (O). The number of assemblages in which a nearest-living relative determined the coexistence interval and/or was categorised as a climate outlier are reported (N) with the total number of assemblages in which it was found reported in brackets.

| Nearest-living relative | N | MAT | CMT | WMT | MAP | HMP | LMP | WMP |
| --- | --- | --- | --- | --- | --- | --- | --- | --- |
| *Family-level* |  |  |  |  |  |  |  |  |
| [P] Araucariaceae | 9 (12) | - | LB/O | - | LB | LB | - | - |
| [P] Cyrillaceae | 6 (9) | - | - | UB | - | LB | - | LB |
| [P] Juglandaceae | 1 (26) | - | - | - | - | - | - | UB |
| [P] Pteridaceae | 7 (31) | - | - | UB | - | LB | - | LB/O |
| [P] Santalaceae | 2 (4) | - | - | UB | - | N | LB | - |
| [P] Schizaeaceae | 10 (10) | UB | UB | UB | LB/O | LB | LB/O | LB |
| [P/I] “taxodioid Cupressaceae” | 7 (38) | - | - | - | UB | UB | UB | - |
| *Genus-level* |  |  |  |  |  |  |  |  |
| [P] *Cryptomeria* | 2 (2) | - | - | - | LB/O | LB/O | - | - |
| [P/I] *Cycadaceae*<sup>a</sup> | 12 (12<sup>b</sup>) | LB/O | LB | LB/O | LB | LB/O | LB | - |
| [I] *Engelhardia*<sup>b</sup> | 24 (25) | LB/UB | LB/UB | LB | LB | LB/O | - | LB |
| [P] *Hemiptelea*<sup>a</sup> | 1 (1) | - | UB | LB | - | N | - | LB |
| [P] *Keteleeria* | 14 (14) | LB | LB | - | LB/O | LB/O | - | LB/O |
| [P] *Liquidambar* | 9 (27) | LB | LB | LB | - | LB | - | - |
| [P] *Pinus*<sup>c</sup> | 1 (41) | - | - | - | - | UB | - | - |
| [P] *Planera*<sup>a</sup> | 26 (26) | LB/UB | LB/UB | LB/UB | LB/UB | LB/UB/O | LB/O | LB/O |
| [P] *Podocarpus* | 13 (34) | LB | LB | - | - | N | LB | LB |
| [P] *Ranunculus* | (7) | - | UB | UB | - | N | - | UB |
| [P] *Rhus* | 36 (40) | - | - | LB | LB/UB | UB/O | LB | LB |
| [P] *Sabal* | 4 (5) | LB | - | LB/O/UB | - | N | - | - |
| [P] *Taxodium* | 1 (2) | - | - | - | UB | N | - | - |

MAT = mean annual temperature; CMT = mean temperature of the coldest month; WMT = mean temperature of the warmest month; MAP = mean annual precipitation; HMP = highest monthly mean precipitation; LMP = lowest monthly mean precipitation; WMP = mean precipitation of the warmest month.

Of the 2207 listed associations (in total, including duplicates; Quan et al., 2012, appendix 1), 1) 115 were flagged as uncertain (e.g. “Osmunda?”); 2) we found 66 associations where it was unclear which systematic concept was used (e.g. “Cycadaceae” may have been used as a synonym for “Cycadales” judged from the reported associations); and, 3) an additional 24 cases cited obsolete taxa (not counting “taxodioid Cupressaceae”). The details regarding invalid, problematic, uncertain (flagged as “?” in Quan et al.) or systematically unclear associations are provided in **File S1** in the online supporting archive (OSA) available for anonymous download at http://www.palaeogrimm.org/data/Grm15FnF_OSA.zip.

There are four problems highlighted by the use of artificial pollen genera by Quan et al. (2012), and thereby must be inherent in the Palaeoflora database. Firstly, artificial pollen genera are associated with modern genera that have no diagnostic pollen at all, or pollen easily confused with other related or unrelated modern genera (see **File S1**). Secondly, in many cases the chosen nearest-living relative at the genus-level has tolerances that are not representative for the entire group of taxa that may equally be considered nearest-living relatives of the fossil taxon; this typically results into too narrow intervals (**Figs 2, 3**; see **File S2** in OSA for complete list).

**Figure 3.**
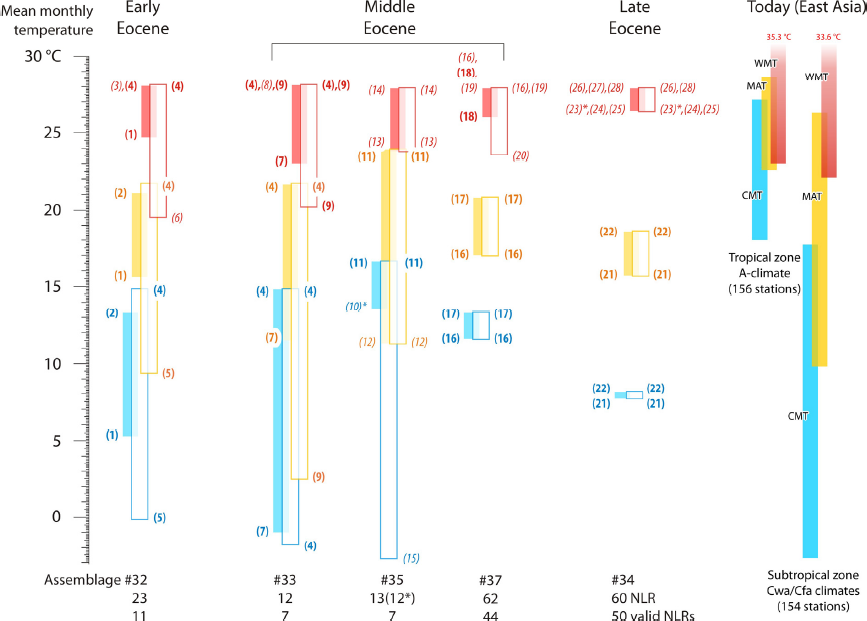
Reconstructed ‘palaeotemperatures’ for the Eocene of southernmost China using the Coexistence Approach and the Palaeoflora database. The CMT, MAT, and WMT intervals as a) reported in Quan et al. (2012, table 2) (solid boxes), and b) after elimination of duplicate, uncertain, problematic and invalid NLRs and obviously erroneous tolerances (open boxes). Nearest-living relatives (NLR) defining coexistence intervals are indicated by Arabic numbers; in italics when informing a single interval boundary, in bold font when informing two or more interval boundaries: (1) *Engelhardia*, wrong NLR; (2) *Planera*, highly problematic NLR; (3) Loranthaceae, uncertain NLR; (4) *Persicaria–Polygonum*; (5) *Ephedra*; (6) Olacaceae; (7) *Liquidambar*, problematic NLR; (8) *Celtis*; (9) *Ostrya*; (10) Sonneratiaceae, invalid NLR (* although not recognised as ‘climatic outlier’ in this particular assemblage, MATmin and WMTmin of Sonneratiaceae were apparently not considered by Quan et al. to determine coexistence intervals); (11) Polypodiaceae; (12) *Eurya*; (13) Lygodiaceae (=*Lygodium*); (14) *Nyssa*; (15) Proteaceae; (16) *Ocotea* (Lauraceae of the Americas, with a few species in Africa; reported as macrofossil); (17) *Tilia* (reported as microfossil); (18) *Sabal*, problematic NLR; (19) Podocarpaceae (cosmopolitan family); (20) Symplocaceae; (21) *Trema*; (22) *Corylopsis*; (23) *Macaranga* (* tropical genus, recognised as ‘climatic outlier’ for MAT and CMT intervals); (24) *Lotus*; (25) *Skimmia*; (26) Onagraceae; (27) *Ranunculus*, problematic NLR; (28) *Typha*.

Thirdly, as highlighted above, pollen with an ambiguous generic affinity (**Table 5**) is sometimes represented by artificially combined nearest-living relatives (see also Grimm and Denk, 2012). In all but one of the cases with artificially fused nearest-living relatives, the tolerance range is the coexistence interval of both unrelated taxa. For example, *Boehlensipollis,* according to Quan et al. (2012, appendix A), corresponds to either Sapindaceae (Sapindales; a wrong association) or Loranthaceae (Santalales, a possible association; Huang, 1972). The climate range of *Boehlensipollis* is estimated in Quan et al. (2012) as the mutual climate range *shared* by Sapindaceae and Loranthaceae (i.e. the overlap in climate space, not the total range of climate space, between these families). This is in direct violation of Coexistence Approach assumptions (1 and 2) and the nearest-living relative principle. If researchers wish to include taxa with ambiguous affinities in an MCR-NLR method, the entire climate tolerance realised by all modern taxa belonging to the same phylogenetic lineage need to be considered in analogy to the use of genus- and family-level nearest-living relatives for fossils with ambiguous species/genus affinities. In the case of *Boehlensipollis*, this would be the entire rosid clade (**Table 5**), which includes *all descendants of the last common ancestor* of Loranthaceae (cf. Huang, 1972), Elaeagnaceae and Lythraceae (e.g. *B. hohlii*; Stuchlik et al., 2014). The rosid clade is cosmopolitan and would provide a wide and meaningless set of climate tolerances. The current illogical treatment of ambiguous fossil taxa however generates tolerance ranges that are too narrow, biologically artificial, and simply erroneous; this increases the likelihood of reconstructing narrow and pseudo-precise climate intervals (see assemblage #63, **Fig. 2**).

**Table 5.**
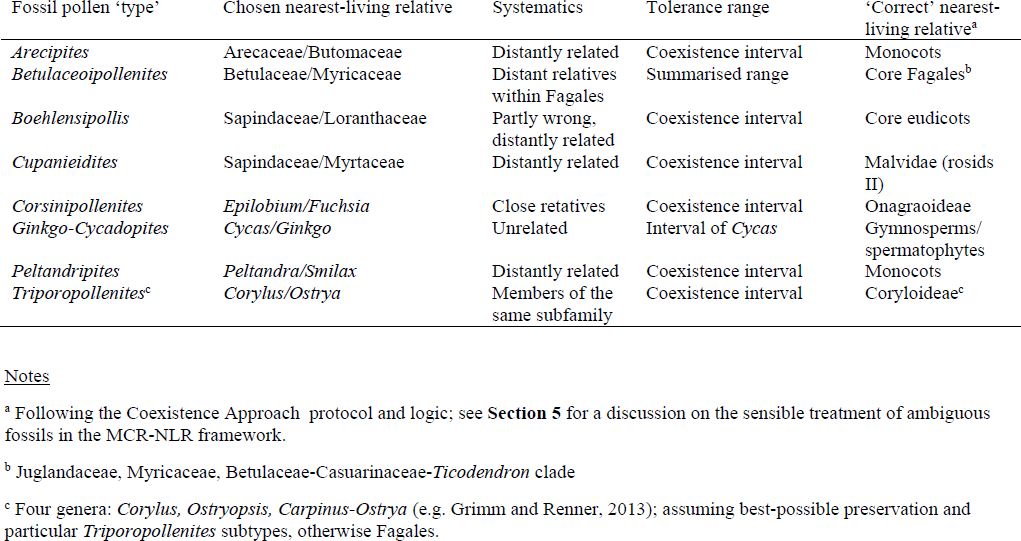
Examples of erroneous nearest-living relatives reported in Quan et al. (2012) that comprise artificially-joined taxa when the fossil pollen type was ambiguous. In most cases the coexistence interval (the mutually shared climate range) of two, often distantly related taxa was chosen; a direct violation of assumptions 2 and 3 of the Coexistence Approach.

The last problem highlighted is that poorly preserved or indistinct pollen are represented either by family-level nearest-living relatives or by using a specific genus and family but adding a “?”. Although never explicitly stated by Quan et al. (2012) how these uncertain nearest-living relative associations were included in the analysis, it is clear that they were used unconditionally, occasionally defining intervals, forcing the recognition of ‘climatic outliers’ or triggering ‘ambiguous’ intervals (Quan et al, 2012: table 2 versus appendix B; **File S2** in OSA). Quan et al. also do not report the number of ‘outliers’ or ‘ambiguous’ intervals, although they could be reconstructed based on the data provided in appendices A and B of that study.

We have highlighted a number of problematic nearest-living relatives used in the study of Quan et al. (2012) that define climate intervals (**Table 4**) and this illustrates the inherent “pitfalls” (Grimm and Denk, 2012) and “uncertainties” (Utescher et al., 2014) of CA+PF studies. When invalid, problematic or uncertain nearest-living relatives are eliminated, reconstructed climate intervals tend to be far less resolved (**Fig. 3**).

##### Box 1. The case of *Engelhardia* and ‘*Cupuliferoipollenites*’

Coexistence Approach studies often use wrong or non-representative nearest-living relatives. For example, Utescher et al. (2014, p. 63) state that *Engelhardia* is a reliable proxy for palaeoclimate reconstruction. However, they overlook that all fossils (leaves, fruits, and pollen) from the northern hemisphere that have been referred to as ‘*Engelhardia*’ are actually members of the Engelhardioideae, a subfamily of the Juglandaceae including four modern genera (Jähnichen et al., 1977; Manchester, 1987; Kvaček, 2007; Manos et al., 2007). The majority of fossil leaves most closely resemble the foliage of the modern genus *Oreomunnea*, one of two currently accepted New World genera with two geographically restricted species in Mexico/Central America. Two of the three western Eurasian fossil taxa, based on reproductive structures, are intermediate between the modern East Asian *Alfaropsis* and the New World *Oreomunnea*, and the last resembles *Alfaropsis*; therefore Jähnichen et al. (1977) suggested that the fossil taxon represent an extinct lineage of Engelhardioideae. Pollen of all four extant genera of the Engelhardioideae are highly similar, even when investigated under the scanning electron microscope (Stone and Broome, 1975). Pollen indistinguishable from extant Engelhardioideae has also been reported for other extinct Juglandaceae (Manchester, 1987). Hence, the nearest-living relative for such fossil pollen can be, at best, Engelhardioideae and certainly not *Engelhardia*, which is a genus currently restricted to tropical Southeast Asia, with two geographically restricted species extending into subtropical southern China. The most widespread and cold-tolerant of the four extant genera is the monotypic East Asian *Alfaropsis* (= *Engelhardia roxburghiana* in Flora of China). The temperature tolerances recorded in \the Palaeoflora database of *Engelhardia* do not include the climatic range of the more cold-tolerant *Alfaropsis (Engelhardia) roxburghiana*. Initially, a minimum MAT tolerance (MATmin) of 17.5 °C was used for *Engelhardia* (Mosbrugger and Utescher, 1997). This value was later changed to 15.6 °C (Quan et al., 2013 and all Coexistence Approach literature that documented climate-delimiting taxa), and eventually to 13.6 °C (Utescher et al., 2014, table 2). The latest updated estimate is still 2–3 °C higher than the conservative MATmin estimate for Engelhardioideae of 10–11 °C (Fang et al., 2009, corrected for altitudinal bias; Grimm and Denk, 2012). Moreover, based on the fossil record it is highly unlikely that no extinct or ancient Engelhardioideae thrived outside the climate niches of the modern, disjunct genera. Hence, Quan et al.’s (2013) use of *Engelhardia* for pollen with affinity to Engelhardioideae (or extinct Juglandaceae) violates three Coexistence Approach assumptions: wrong selection of nearest-living relative (violation of assumption 1, main-text **Table 2**), climatic niche of the fossil taxon is different from extant taxa (violation of assumption 2, main-text **Table 2**), and potential climate niche can only be estimated for *Alfaropsis roxburghiana* based on known distribution (partial violation of assumption 3, main-text **Table 2**). In Quan et al. (2013), the erroneous nearest-living relative ‘*Engelhardia*’ defines the lower boundaries of MAT, CMT *and* WMT intervals (simultaneously) for eight assemblages, of MAP in one and WMP in two assemblages (main-text **Table 3**; in assemblage #63, five of seven lower boundaries are defined by *Engelhardia*, **File S2** in the online supporting archive), including two from southern China which are fundamental for the authors’ conclusions (Quan et al., 2013, fig. 4). Thus, this nearest-living relative was critical in many of the Coexistence Approach climate reconstructions of Quan et al. (2013). If *Engelhardia* would be excluded (or replaced by Engelhardioideae), CMT values would become largely uninformative, MAT values would cover temperate to tropical conditions, and only the reconstructed WMT intervals would remain indicative for hot summers. However, these ‘better’ intervals would, in turn, rely on other problematic nearest-living relatives (main-text **Fig. 2**).

Identification of nearest-living relatives from pollen is not a trivial endeavour and the “*Cupuliferoipollenites*” offer a further example of this. “*Cupuliferoipollenites*” is generally used for pollen grains with highly similar surfaces under light microscopy and morphological affinity to modern Fagaceae; this includes extinct genera such as *Trigonobalanopsis* which can only be differentiated using scanning electron microscopy from modern members of the Castaneoideae (*Lithocarpus, Chrysolepis, Notholithocarpus, Castanea, Castanopsis*) and the trigonobalanoids (*Trigonobalanus*; pollen of *Colombobalanus* and *Formanodendron* are different). Pollen of the modern Castaneoideae, are barely distinguishable under the scanning electron microscope (e.g. Bouchal et al., 2014; Grímsson et al., 2014a) despite the substantial genetic differentiation between the modern genera and complex phylogenetic relationships, particular with respect to the largest Fagaceae genus *Quercus*, the oaks (Manos et al., 2001; Manos et al., 2008; Denk and Grimm, 2010; Hubert et al., 2014). One can assume that no Fagaceae lineage thrived outside the combined climate space of *Lithocarpus and Quercus*; *Quercus* pollen can be unambiguously identified under the light microscope, and therefore the cumulative climate space of *Lithocarpus–Castanea/Chrysolepis* (as the most tolerant genera) may be used to circumscribe the climate space of *Cupuliferoipollenites*. However, as *Quercus* nests deep within the Fagaceae in various molecular trees, it cannot be discounted that an extinct sublineage had a climatic niche more similar to the one today covered by *Quercus*. Hence, the nearest-living relative for *Cupuliferoipollenites* pollen would be Fagaceae excluding *Fagus* (i.e. the cumulative range of *Lithocarpus* and *Quercus*). Unaware of current taxonomy and Fagaceae systematics, Quan et al. (2013) used tolerances that equal exactly those recorded for *Castanea*, a small but widespread genus, not extending into the warm subtropics or tropics, inflicting minimum tolerance errors – based on Palaeoflora data –– of up to 10 °C for temperature, 1200 mm/year, or 180 mm/month for precipitation (**File S2**). In contrast to the use of *Engelhardia* as nearest-living relative for all Engelhardioideae, these massive errors had no effect on the reconstructed intervals as all Fagaceae rarely interfere with the Coexistence Approach results due to their relatively broad climatic tolerance. However, it does provide an apt example of the problem of identifying nearest-living relatives for fossil taxa. Each and every association requires extensive consideration, investigation and reporting, else incorrect assignment can easily lead to substantial biases in Coexistence Approach findings.

## 3 Determination of minimum and maximum tolerances

### 3.1 Tolerance data in the Palaeoflora database

Grimm and Denk (2012) noted that the frequent reason for highly-resolved and pseudo-precise MAT intervals in CA+PF studies was due to erroneous nearest-living relatives and/or tolerance values in the Palaeoflora database that were too narrow (see also **Figs 2, 3**; **File S2**). They also offered a number of examples that raised doubts about the representativeness of MAP tolerance data as indicated in the Palaeoflora database. The documentation published by Quan et al. (2012) provides a rare opportunity to further explore these concerns, particularly regarding other reconstructed climate parameters. **Figure 4** shows the reported minimum and maximum tolerances of nearest-living relatives and Coexistence Approach intervals for the richest assemblage with a total of 102 recorded, distinct nearest-living relatives (for a flora of 179 distinct fossil taxa: 176 palynomorphs, 3 macrofossils). The following can be observed: *(i)* Only few nearest-living relatives have minimum or maximum tolerances similar to those that actually determine the intervals. *(ii)* Comparatively high to very high values are evident for the medians of the mean temperature of the coldest month (CMT) and mean annual temperature (MAT) – less than 4 and 2 °C lower than the median of the mean temperature of the warmest month (WMT) max-tolerances. *(iii)* Precipitation tolerances for the warmest month (WMP) are substantially *lower* than for the month with the highest precipitation (HMP) – despite most nearest-living relatives thriving today in the warm temperate climates of (southern) China characterised by *heavy* summer rainfalls. *(iv)* A few ‘climatic outliers’, including two consistently recognised as such (Sonneratiaceae, invalid family, originally including two unrelated genera; Botrychiaceae, obsolete family, ∼ *Botrychium*) and three occasional classified ‘climatic outliers’ (*Engelhardia* [see above]*, Keteleeria, Planera*; **Table 4**) are found; none of which were generally excluded from the analysis according to Quan et al. (2012, appendices A and B; cf. **Files S1, S2**) but were apparently not considered for the calculation of certain intervals (**Fig. 4**; Quan et al., 2012, table 2). *(v)* Nearest-living relatives recognised as ‘climatic outliers’ for one climate parameter (and thus ignored for that analysis) were used to determine alternative or non-ambiguous (pseudo-)precise intervals for other parameters (Cycadaceae, Gleicheniaceae, *Elythranthe, Larix,* Lycopodiaceae; cf. **Table 4**). *(vi)* Only a few taxa are recorded that tolerate high-precipitation tropical-subtropical settings as found today in northeastern India and Burma in contrast to the remaining 95% of nearest-living relatives – north-eastern India and Burma is a known plant biodiversity hotspot. *(vii)* A number of nearest-living relatives that can tolerate or prefer snow (CMT ≤ −3 °C) and even polar climates (WMT ≤ 10 °C) or desert-steppe environments (HMP ≤ 35 mm/month) are found in ‘coexistence’ with a majority of taxa intolerant to such climate niches. Because of the rare MAP, HMP and LMP combination, the reconstructed climate has no modern analogue, hence, represents an “*extinct climate*” (according Utescher et al., 2014, p. 68f). The reconstruction of an ‘extinct’ climate is an indication of a direct violation of the MCR-NLR principles. However, any combination of four out of six parameters (WMP not considered) would agree with values recorded for stations in *Cfa* and *Cwa* climates (**Table S1** in OSA). So the question may simply be whether the recorded minimum and maximum tolerances in the Palaeoflora database fulfil assumption 4 (**Table 2**) of the Coexistence Approach: the modern climate [and distribution] data are of good quality (discussed below and further in the next section).

**Figure 4.**
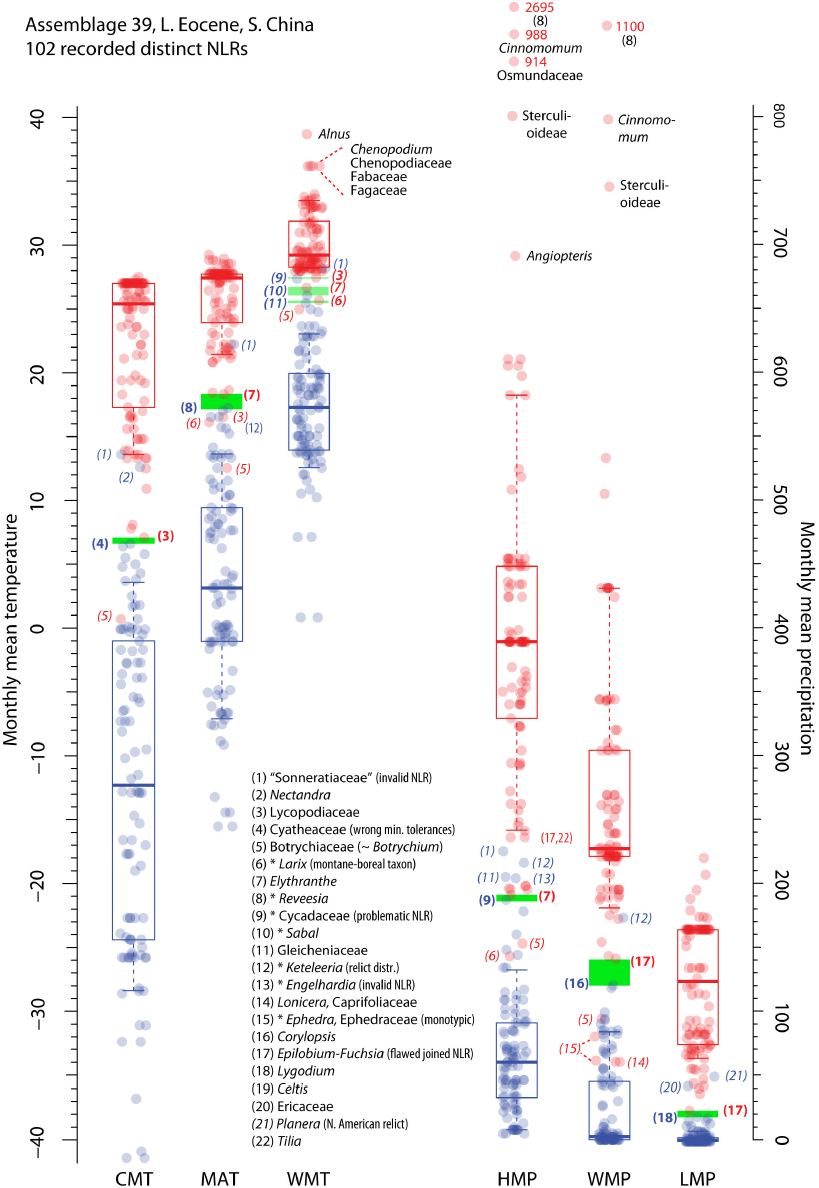
Example demonstrating how coexistence intervals reconstructed by Quan et al. (2012) were directly influenced by exotic (outlier) or problematic nearest-living relatives, even in very rich assemblages. Maximum and minimum climate parameter values for 102 (unique) nearest-living relatives (assemblage 39) are shown in red and blue, respectively. Nearest-living relatives that frequently determine coexistence intervals in other assemblages are highlighted with asterisks. Boxplot whiskers represent 10^th^ and 90^th^ percentiles.

The determination of reasonable tolerance ranges is crucial to the application of the Coexistence Approach or any mutual climate range technique in general (Mosbrugger and Utescher, 1997; Grimm and Denk, 2012; Thompson et al., 2012a; Utescher et al., 2014). The precision (resolution) and accuracy (reliability/reproducibility) of an interval reconstructed for a palaeobotanical assemblage cannot exceed the accuracy at which the tolerances of nearest-living relatives were established. In the case of temperature data, this may be at best an accuracy of 1–2 °C (Grimm and Denk, 2012; Utescher et al., 2014, section 4). Hence a precision (resolution of the reconstruction) of 1 °C cannot be obtained under the current application of the Coexistence Approach, and any increased precision (narrow intervals) relates to accuracy issues. For assemblages constrained by *Engelhardia*, an MAT boundary that is too high by 4– 5 °C (due to a combination of wrong MATmin tolerance + wrong nearest-living relative association) results in a high pseudo-precision of down to 0 °C (assemblage 12 in Quan et al, 2012) for MAT, ∼3 °C for CMT and < 2 °C for WMT (**Fig. 2**). The combination of MAT, CMT and WMT would be indicative of subtropical climates with hot, rainy summers as found today in central (lowlands) and southern China (lowlands and mid-altitudes). The elimination of the erroneous nearest-living relative (*Engelhardia*) results in broader and far less precise estimates. Examples of the new reconstruction precision without this single and problematic nearest living relative include CMT (4–13 °C for assemblages with more than 10 nearest-living relatives) and MAT intervals (3–11 °C in all but one case). Only WMT reconstructions appear relatively stable, leading to MAT, CMT and WMT combinations found in subtropical as well as fully temperate climates of modern China with hot summers.

Further elimination of all problematic nearest-living relatives quickly leads to uninformative climate reconstruction intervals for the majority of these nine floras (not shown, see data provided in **File S2**). Detailed inspection of critical nearest-living relatives (**Table 4**; **Fig. 4**) reveals that phenomena such as narrow climate reconstruction intervals, ‘climatic outliers’, ambiguous intervals, and ‘extinct’ climates are direct consequences of inaccurate or unrepresentative (i.e. too narrow) tolerance data. In the light of the uncertainties regarding the best possible data on species distribution and climate data for establishing tolerances of potential nearest-living relatives (Grimm and Denk, 2012; Thompson et al., 2012a; Utescher et al., 2014), it is clear that currently available tolerance data should not be recorded (or used: see Utescher et al., 2014, p. 66) with a purported precision of 0.1 °C and 1 mm precipitation/month and year as has been reported and used in all previous Coexistence Approach studies. However, there is an additional problem when considering the tolerance data in the Palaeoflora database: the subjective bias introduced by handpicking ‘extreme’ climate stations to obtain the tolerance ranges for each species.

### 3.2 Subjective bias: minimum and maximum tolerances in the Palaeoflora database

Utescher et al. (2014) sanctify the Palaeoflora database strategy of selecting 4–6 “*extreme*” climate stations (Mosbrugger and Utescher, 1997) to define the climate tolerance for each extant taxon. They did note that, for example, there are MAT differences of up to 5 °C between the WorldClim data (Hijmans et al., 2005; tested using different grid-sizes) and their selected station data (Müller and Hennings, 2000). However, they suggest that this discrepancy is due to ‘station data-holes’ and subsequent inaccuracy in the extrapolated data (compare Kottek et al., 2006; and Peel et al., 2007); however, it is more likely to be due to different observation periods of the climate data: a maximum of 30 years in Müller and Hennings (2000) and 50 years for WorldClim interpolations (with the exclusion of stations with incomplete time coverage). Walter and Lieth (1960) illustrated the fluctuations of mean values with climate diagrams for Hohenheim (*Cfb* climate, Central Europe) covering the years 1907–1956; this showed an MAT and MAP variation of between 7.1–10.1 °C (8.4 °C average) and 459–975 (685) mm/year, respectively. In the light of such fluctuations, and contrary to the assertions by Utescher et al. (2014) that highly precise values are methodologically sound, we reiterate that it is absurd to estimate tolerances based on 4–6 ‘extreme’ stations with an accuracy of 0.1 °C and 1 mm precipitation per month or year.

Furthermore, selection of a “*nearby*” climate station in areas with too few climate stations to capture local climate differences is not less error-prone than using an extrapolated climate surface (Grimm and Denk, 2012, ES4). **Figure 5** shows CMT tolerance profiles for North American trees and shrub genera (Thompson et al., 1999b, a, 2001; Thompson et al., 2006; Thompson et al., 2012a) compared to Palaeoflora CMT tolerances reported in Quan et al. (2012).

**Figure 5.**
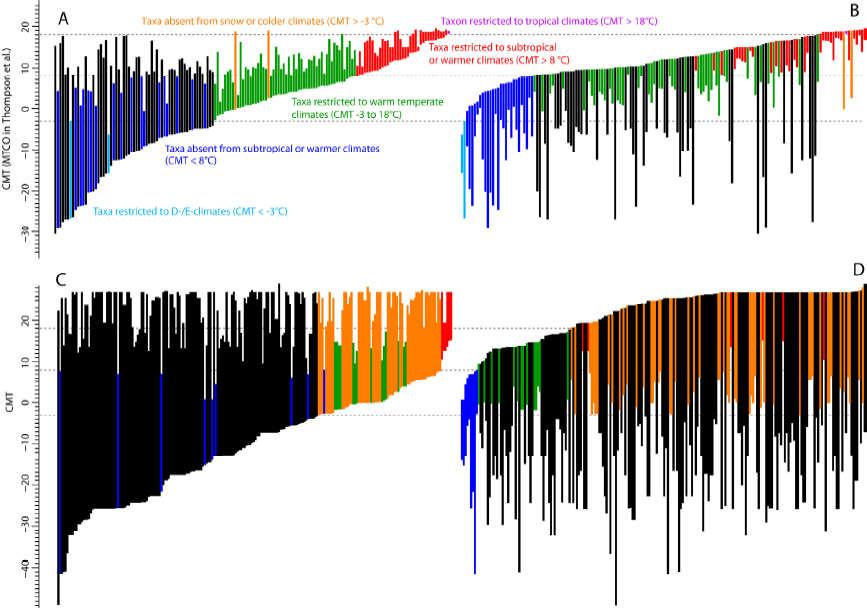
Contrasting tolerance profiles of the mean temperature of the coldest month (CMT) for (A) North American tree and shrub genera (obtained from gridded data; Thompson et al., 2012b) versus (B) nearest-living relatives from the Palaeoflora database (obtained from the selection of 5-6 climate stations). The nearest-living relatives range from genus to family levels.

As expected, the tolerance profiles for North America show an appreciable variance as they are based on relatively well-resolving 25 km-gridded distribution and climate data that only has some limitations for taxa confined to western North America (Thompson et al., 2012a). Such variance should be expected as the distributions of genera vary substantially. On the other hand, the Quan et al. (2012) profile is much more uniform, oddly so when considering the upper interval boundaries (**File S4** in online supporting archive [OSA] provides profiles for all seven parameter). This explains the high medians for maximum tolerances observed in all assemblages (**Fig. 4**; **File S5** in OSA). In fact, the maximum CMT tolerance values for 77 out of the 246 nearest-living relatives listed in Quan et al.’s appendix B fall within the range of 27–27.5 °C. This is surprising since only five (fully tropical) climate stations out of over 2600 from the northern hemispheric reach such extreme values (see **File S3** in OSA). Tolerance profiles for other parameters show the same deficits and oddities (**File S4**; cf. **Fig. 4**). Obviously, the careful selection of 4–6 ‘extreme’ climate stations fails to capture the substantial differences in the distribution and climate ranges of potential nearest-living relatives (e.g. Fang et al., 2009) and introduces a range of selection biases. Tolerance values generated from gridded data are unlikely to be affected by such selection bias (e.g. Thompson 2001-2006; **Fig. 5A** vs **5B**).

Utescher et al. (2014) explain that temperature and precipitation tolerances are only recorded with a precision of 0.1 °C, and 1 mm/year or month, in the Palaeoflora database to be able to identify the climate station(s) that have been selected (sic!), and not because they believe the tolerance data have such high accuracy. The high redundancy of values in the Quang et al. (2012) tolerance data demonstrate that only a limited set of climate stations is used. Hence, the selection of 4–6 ‘extreme’ stations strongly biases the dataset towards certain values and coexistence intervals (**Fig. 4, File S2, Table S2**) and precludes the possibility to reconstruct fine-scale climate shifts (e.g. 1 °C, 100 mm/year, 10 mm/month).

### 3.3 Quality of data in the Palaeoflora database

The points raised above lead us to question the quality of the tolerance data in the Palaeoflora database. The trustworthiness of the Coexistence Approach is directly dependent on the reliability of the niche characterisation, specifically the tolerances, of the nearest-living-relative taxa. There has been no comprehensive assessment of the reliability of tolerance data, other than MAT (Grimm and Denk, 2012) in the Palaeoflora database (assumption 4 of CA; **Table 2**), and it is beyond the scope of this study to quantify the errors in the data provided by Quan et al. (2012) for all seven climate parameters. It is clearly up to the curators and applicants of Palaeoflora data to demonstrate that their data comply with the four assumptions of the Coexistence Approach (**Table 2**). However, we will point out some of the most obvious errors and problems. First, the systematic uncertainties inherent to the Palaeoflora data, both regarding the association of fossils to nearest-living relatives and the circumscription of important, interval-defining higher level nearest-living relatives (such as “taxodioid Cupressaceae”, *Engelhardia*, Cycadaceae; **Table 4**) require a constant and documented curating effort. Family-level intervals should always be greater or equal to the intervals of the constituent genera or species, and never the other way round. **Table 6** shows that even in the case of monotypic families, such inconsistencies are found between the family and genus tolerances. These inconsistencies affect reconstructed intervals (highlighted in **Table 6**). Using the dataset provided by Quan et al. (2012), we looked for inconsistencies in 38 family and subfamily tolerances recorded in the primary Palaeoflora database using the recorded tolerances of their constituent genera. We detected inconsistencies (either too wide genus tolerances or too narrow family tolerances) of up to 14 °C in MAT and WMT (e.g. monotypic Selaginellaceae), 24 °C in CMT (e.g. Caprifoliaceae, family and three out of c. 42 genera listed), 7600 mm/year in MAP, 2300 mm/month in HMP, and 800 mm/month in WMP (e.g. Malvaceae, family and 6/243 genera listed), and 90 mm/month in LMP (e.g. Polypodiaceae). Likely causes for such alarming inconsistences are

1. poor systematic control (Grimm and Denk, 2012; Utescher et al., 2014). The Pinaceae is the only family for which the recorded family tolerance corresponds to the subsumed tolerances of all genera;
2. the strategy of the database curators to only check Palaeoflora entries if the nearest-living relative is frequently recognised as ‘climatic outlier’ (Mosbrugger and Utescher, 1997; Utescher et al., 2014);
3. the frequent usage of the interval of one genus to define the tolerance of an entire family: Appendix B to Quan et al. lists 12 non-monotypic, partly highly diverse families with intervals identical to that of a single genus;
4. unavoidable subjective errors during the careful selection of 4–6 ‘extreme’ climate stations (Utescher et al., 2014, p. 66); and
5. bulk definition of tolerances, in particular of maximum values, for taxa that do not affect Coexistence Approach reconstructions (cf. **Fig. 5B**). For instance, one fifth (43 out of 219) of nearest-living relatives listed by Quan et al. have the ***same*** MATmax = 27.7 °C, CMTmax = 27 °C, MAPmax = 3151 mm/year, HMPmax = 389 mm/month, and LMPmax = 165 mm/month tolerances.

**Table 6.** Examples of inconsistencies (> 1°C, >100 mm/year, 10 mm/month) in the Palaeoflora database where family-level tolerances do not correspond to genera-level tolerances. These examples affected the climate reconstructions of Quan et al. (2012). The minimum error is defined by how much genus tolerance(s) exceed the tolerance recorded for its (their) subfamily and family. See **File S2** in online supporting archive for the full list including 38 family- or subfamily-level taxa.

| Family/subfamily | Total number of genera | Genera listed in Quan et al. (2012) | Parameter/value | Minimum error |
| --- | --- | --- | --- | --- |
| Taxodioideae | 4 | 3 | MAP max | 600 mm/year |
|  |  |  | HMP max | 200 mm/month |
|  |  |  | LMP max | 40 mm/month |
| Cycadaceae | 1 | 1 | WMT min | 4 $^{\circ}\text{C}$ |
|  |  |  | MAP min | 100 mm/year |
| Lygodiaceae | 1 | 1 | MAP max | 200 mm/year |
| Polypodiaceae | c. 40/50 | 1 | MAT max | 5 $^{\circ}\text{C}$ |
| | | | CMT max | 10 $^{\circ}\text{C}$ |
|  |  |  | LMP max | 90 mm/month |
| Pteridaceae | c. 40/50 | 2 | WMT max | 4 $^{\circ}\text{C}$ |
|  |  |  | WMP min | 30 mm/month |
| Santalaceae | 44 | 1 | WMT max | 5 $^{\circ}\text{C}$ |

Judged from the data released to the public so far, the widely used Palaeoflora database (Utescher and Mosbrugger, 2009) does not (yet) fulfil assumption 4 of the Coexistence Approach (Grimm and Denk, 2012) as acknowledged by most recent studies applying this method (Eldrett et al., 2014; Kotthoff et al., 2014). Utescher et al. (2014) provide a comprehensive review on the reliability of modern climatic data and conclude correctly that “*it is difficult to quantify predictive uncertainty. A few degrees difference regarding temperature, and some tens of millimetres with respect to annual precipitation can be cited as a rough assessment (cf. also Mosbrugger and Utescher, 1997), … Even with high resolution chorological data, greater precision can hardly ever be achieved due to uncertainties in the meteorological observations, be they station data or derived from gridded datasets.*” A few (≤ 3 °C) degrees uncertainty regarding temperature covers most climate shifts detected and discussed in Coexistence Approach literature, including the Quan et al. study (**Figs 2, 3**; **File S5**).

### 3.4 What is a relict, and at which point does it become a problem for the Coexistence Approach?

An additional problem that is not addressed within in the Coexistence Approach framework, but is nonetheless a crucial issue, is that of relictual distributions of identified nearest-living relatives. Taxa for which the modern distribution is restricted and cannot be considered representative of their distribution in the past (assumption 2 of CA, **Table 2**) include Engelhardioideae (Manchester, 1987; Kvaček, 2007; Fang et al., 2009; Flora of China, 2014), *Taxodium* (Thompson et al., 1999b), and *Keteleeria* (Fang et al., 2009; Flora of China, 2014). One has to keep in mind that the distribution of a plant never is solely controlled by climate, but the result of other modern and historical biotic and abiotic factors (Grimm and Denk, 2012; Utescher et al., 2014). The critical question is how much of the potential climatic niche of the taxon is actually realized by the taxon (3rd assumption of Coexistence Approach). However, in many Coexistence Approach studies, such as Quan et al. (2012; **Table 2, Figs 2–4**), exactly those taxa that are problematic regarding the 2nd and 3rd assumption define the intervals (Grimm and Denk, 2012, ES2), whereas unproblematic taxa have reported tolerances that are usually too broad to be informative in the Coexistence Approach framework. As consequence, substantial differences between reconstructed climate intervals are not too frequent in CA+PF studies, at least for MAT values (Grimm and Denk, 2012; Utescher et al., 2014, fig. 7). This is also illustrated in **File S5** for Coexistence Approach versus statistically-controlled mutual climate range reconstructions for all fossil assemblages compiled by Quan et al. (2012). If taxa with small distribution areas are excluded, any differences in the reconstructed intervals fade out even further.

## 4 Unsettling phenomena in the application of the Coexistence Approach

### 4.1 Examples of informative nearest-living-relatives used for the Eocene of China that violate the basic assumptions (Table 2) of the Coexistence Approach

The accuracy of the Coexistence Approach relies upon thoughtful, investigative and *documented* consideration of each association of the fossil taxon and a nearest-living relative. However, we suspect that the majority, if not all, CA+PF studies fall under the adage of ‘garbage in, gospel out’. Here we restrict ourselves to reviewing examples of ‘informative’ nearest-living relatives – i.e. those responsible for establishing the CA+PF climate reconstruction for a given assemblage – in the Quan et al. (2012) study.

**Cycadaceae/*Cycas*** are among the most important nearest-living relatives of the study of Quan et al. (2012). These taxa constrain reconstructions to relatively high values of MAT (≥ 16.5 °C; **Fig. 4**), CMT (≥ 5.5 °C) and WMT (≥ 27.3 °C, Cycadaceae only) as well as high values for MAP (≥ 887 mm/year), HMP (≥ 187 mm/month), and LMP (≥ 8 mm/month; i.e. no month without rainfall). This generated a number of ambiguous intervals and the taxon was occasional eliminated as an outlier (**Table 4**). The family is monotypic – *Cycas* is the only genus – and the lineage probably diverged from the rest of the Cycadales in the late Palaeozoic/early Mesozoic (Gao and Thomas, 1989a, b). Nevertheless, pollen of *Cycas* (Cycadaceae) and *Encephalartos* (Zamiaceae), both widespread genera, cannot be distinguished under light microscopy (violation of assumption 1). Molecular dating approaches have indicated a relatively young crown age for all Cycadales genera, which would mean that most, if not all, Eocene Cycadales represent earlier radiations (extinct lineages). Therefore the use of the modern distribution (and climate niches) of *Cycas* (or *Encephalartos*) for fossil Cycadales pollen is highly questionable (violation of assumptions 2 and 3). Also, inconsistencies within the Palaeoflora database were detected for the tolerance limits for the Cycadaceae/*Cycas* nearest-living relatives. Family-level tolerance values should span that of all the genera. But the family is recorded to be 4 °C less tolerant for WMT than the genus *Cycas* (**Table 6**), and this error resulted in six potentially wrong and four ‘ambiguous’ intervals (violation of assumption 4).

In the case of **Cyatheaceae**, a highly diverse group of tree ferns, the association between pollen and nearest-living relatives is unproblematic. *Cyathidites/* Cyatheaceae pollen is recorded for 15 assemblages from all three reconstructed “*climate zones*” (according Quan et al., 2012, appendix A) – “*I, humid warm temperate to subtropical*”, “*II, middle arid (subtropical heights)*”, “*III, tropical to subtropical*” – and time slices – early, middle, and late Eocene (e.g. **Fig. 4**). This was unsurprising regarding the modern tropical to southern temperate distribution and diversity of the family. What was cause for alarm were the tolerances recorded for such a taxonomically and climatically diverse family. For example, *Cyathea* grows on Steward Island, an island south of New Zealand. Comparing the values of the closest climate station to this island (Ivercargill Airport) with the Palaeoflora database shows a range of inconsistencies: MAT_min_ (∼10°C versus 15.2°C, respectively), CMT_min_ (∼5°C versus 6.6 °C) and WMT_min_ (∼14°C versus 19.6°C); this is a violation of assumption 4. Using these values, this family would become uninformative for MAT reconstructions (**Table 4**); in case of CMT, other problematic nearest-living relatives would dictate the lower boundaries (**Table S3** in OSA).

***Planera*** (**Table 4**, see also **Figs 2–4**), the ‘water elm’, is a monotypic genus today restricted to southeastern United States. It was used as nearest-living relative for *Ulmoideipites*. This association is a gross and erroneous simplification of the complexity present in this pollen genus. *Ulmoideipites* includes several morphotypes with affinity to different members of the Ulmaceae, as well as forms lacking sufficient diagnostic features to link them to a particular genus (Takahashi, 1989; Jones et al., 1995). *Ulmus,* the largest and most widespread genus of Ulmaceae, can have pollen indistinct from that of smaller genera such as *Hemiptelea* and ***Planera!*** This association of the Eocene pollen with *Planera* is a violation of assumption 1. Moreover, the present-day distribution of the single species of *Planera* is restricted to a small area; it is hence unlikely that the modern distribution is sufficient to estimate climate tolerances of the genus in the Eocene (violation of the 2^nd^ and 3^rd^ assumptions). Following the Coexistence Approach guidelines (Mosbrugger and Utescher, 1997), this monotypic nearest-living relative should have been excluded *a priori*. Instead it defined the lower ***or*** upper boundaries for MAT, CMT, MAP, HMP intervals, and lower boundaries for WMT, LMP, and WMP intervals in ***one third*** of the listed assemblages; occasionally this nearest-living relative generated ambiguous intervals and sometimes was eliminated as a climatic outlier.

***Rhus*** (**Table 4**) is used as nearest-living relative of pollen referred to as *Rhoipites*. Pollen of this type was compared to one species of *Rhus* by Wodehouse (1933), but this association is highly problematic, if not wrong (e.g. Pocknall and Crosbie, 1988). In general, generic affinities of Anacardiaceae pollen are difficult to establish using light microscopy. The family comprises about 80 genera, spanning tropical to temperate climates (Stevens, 2001 onwards). Pollen produced by *Rhus* species cannot be unambiguously assigned to this genus (violation of assumption 1). As a designated nearest-living relative, *Rhus* contributed to the reconstructed temperature and precipitation intervals in 36 out of 40 assemblages in which it was recorded (**Table 4**). In the case of HMP reconstructions, *Rhus* was either recognised as climate-delimiting taxon or ‘climatic outlier’ with no accompanying justification for the choice.

**“Taxodiaceae”** pollen are frequently reported from Cenozoic strata. With the advent of molecular data, the traditionally recognised family has been fused with the Cupressaceae (s.str.), a cosmopolitan conifer lineage comprising many endemic and geographically restricted genera (Earle, 2010). Quan et al.’s (2012) fossil lists include a total of seven taxa (**Table S4**), and use a nearest-living relative labelled as “taxodioid Cupressaceae” for pollen ‘typified’ by artificial pollen genera in addition to the three modern genera *Cryptomeria, Glyptostrobus,* and *Taxodium*; three genera that form the current subfamily Taxodioideae (Earle, 2010). However, pollen of these three genera cannot be distinguished using light microscopy and only under using scanning electron microscopy if studied in great detail (Plate I; violation of assumption 1). “Taxodioid” fossils were not only much more widespread in the past (e.g. Manum, 1962), but also probably more diverse. Hence, it is unlikely that the climatic niche of their few survivors, two species in East Asia, *Cryptomeria japonica* and *Glyptostrobus pensilis*, the latter listed as “endangered”, and the species in North and Middle America (*Taxodium distichum*, which likely includes *T. mucronatum*; Earle, 2010, http://www.conifers.org/cu/Taxodium.php, accessed 6/11/2014) can inform the potential or actual climate range of the Eocene Chinese members of the subfamily (violation of assumption 2). The highly restricted modern range of all three (or four) species (Thompson et al., 1999b; Flora of North America, 2004 onwards; Fang et al., 2009; Earle, 2010; Flora of China, 2014) barely represents the potential range of the Taxodioideae in the recent or distant past (violation of assumption 3). Here again we found inconsistencies in the tolerance data documented in Quan et al. (2012): the genus tolerances of two of the three monotypic genera are much wider than the tolerance recorded for the higher order nearest-living relative “taxodioid Cupressaceae”, which should incorporate them, and differ from other sources of bioclimatic data (violation of assumption 4). Even though ***all*** basic assumptions are violated, the “taxodioid Cupressaceae” nearest-living relative was not removed from the analyses (**Table 4, File S2**) contrary to the guidelines of the Coexistence Approach inventors (Mosbrugger and Utescher, 1997; Utescher et al., 2014).

### 4.2 Chronic instability and inconclusiveness of CA+PF reconstructions, and a brief introduction to alternative MCR-NLR techniques

Eocene assemblages of China have different reconstructed climates using the Coexistence Approach almost entirely due to a small minority of nearest-living relatives that deviate in their tolerances from the rest of the assemblage (**File S2**). The absence or presence of any nearest-living relatives from this minority can dramatically change the reconstructed interval. However, this is not the case for MCR techniques in general (Thompson et al., 2012a).

There are several simple means to counter distorting effects of exotic or outlier tolerance values: (*i*) **‘capped’ MCR** (or CA) does not use minimum and maximum tolerances but the 10^th^–90^th^ or 25^th^–75^th^ percentiles. This strategy was recommended by Grimm and Denk (2012) for cases where high quality distribution data are available, such as those provided by Thompson et al. (1999-2012). The reasoning behind ‘capped’ MCR is simple: plants usually do not thrive at the margins of their climatic niches. Hence, the probability is quite low of finding two plant taxa preserved as fossils, both close to their minimum or maximum tolerance. ‘Capped’ MCR can be seen as a simplification of the ‘weighted’ MCR technique proposed by Thompson et al. (2012a). We expect that ‘capped’ MCR will outperform any other MCR-NLR strategy in the identification of mixed palynofloras, such as assemblages with allochtonous elements representing different climate zones. Note that this approach cannot be used if the underlying tolerance data is generated from a handful of climate stations, as is the case for the Palaeoflora database.

(*ii*) **Statistically-controlled (SC-)MCR** identifies statistical outliers in the set of nearest-living relatives, i.e. tolerance values that are substantially different from the values recorded in the rest of the flora, which then are eliminated for the reconstruction of the coexistence interval (Greenwood et al., 2005). The thresholds selected for such statistical controls are commonly based on two-times the standard deviation. However, the number of nearest-living relatives that can be identified for a fossil flora is typically too limited to use this expansive threshold criterion. Therefore, the threshold criterion should be based on absolute numbers. SC-MCR using the 10^th^ (for maximum tolerances) and 90^th^ (for minimum tolerances) percentiles has recently been applied in two studies focussing on North America (Eldrett et al., 2014; Kotthoff et al., 2014) to avoid the problems of encountered in the CA+PF highlighted by Grimm and Denk (2012). By using the percentile cut-off, SC-MCR will eliminate (most) ‘climatic outliers’ as well as ‘ambiguous’ intervals. SC-MCR should not be confused with ‘capped’ MCR/weighted MCR as they apply the statistical filtering at different analytical steps (e.g. Eldrett et al., 2014; Kotthoff et al., 2014).

### 4.3 Do all roads still lead to North Carolina?

Grimm and Denk (2012) noted that most climate parameter combinations found in the Coexistence Approach literature have reconstructed MAT, CMT, WMT and MAP values, which are today found in the subtropical, per-humid lowland *Cfa* climate of North Carolina. Using MAT values of over 500 taxa recorded in the Palaeoflora database, it was apparent that most of these nearest-living relatives can co-exist under such a climate. Of the 50 “*subtropical*” and “*temperate*” elements included in the original *“significance test”* by Mosbrugger and Utescher (1997) to establish the minimum coexistence percentage for a “*statistically significant”* Coexistence Approach interval, 48 can co-exist with all others (notably including taxa with recorded tolerances that are too narrow). However, Utescher et al. (2014, p. 70) suggested that CA+PF is not fundamentally flawed and that – despite the many sources of error – it represents a robust framework that is a “*significant and versatile tool*” for palaeoclimate reconstruction. They are also convinced that with increasing number of nearest-living relatives, one will obtain more precise and accurate reconstructions.

As outlined above, the Palaeoflora database tolerance profiles appear artificial for all recorded parameters (**Fig. 5**; **File S4**). There are very few taxa with tolerances that could potentially reconstruct fully temperate or tropical conditions, or exclude warm, subtropical (CMT ≥ 8 °C) *Cwa*/*Cfa* climates. Furthermore, the maximum number of possibly informative intervals, i.e. non-overlapping intervals, is strongly limited in this list (**Table S2**). For instance, potentially tropical conditions (CMT ≥ 18 °C) or cold/warm summers (WMT ≤ 22 °C) are beyond the resolution of the recorded Palaeoflora nearest-living relatives tolerances as well as high-precipitation regimes found within todays *Aw/Cwa* climates because of the East Asian summer monsoon. Resolving different conditions realised within the warm *Cwa/Cfa* climates of (modern) China showing MAP ranging between 870–2800 mm/year and MAT of 17–24 °C (data from 27 stations, Yunnan–Fujian) is strongly limited based on these tolerance data. The inability to reconstruct tropical climates is particularly surprising given that the set of nearest-living relative taxa covers Eocene assemblages that grew at low latitudes during a global climate optimum, hence, on would expect a substantial and distinct tropical element/signal in those floras. If all data provided by Quan et al. are pooled, one reaches 94% ‘coexistence’ for a hypothetical climate with MAT of 17 °C (208 of 219 NLR, 94% ‘coexistence’), CMT of 7.2 °C (206/219; 94%), WMT of 25.5 °C (210/219; 96%), and MAP of 1215 mm/year (212/219; 97%), very similar climates are found in Georgia, southeastern USA (stations Newnan 4NE, Talbotton 1NE, Washington 2ESE, WMO, station codes: 7221902, 7221707 and 7231105). Coexistence intervals reconstructed for Quan et al.’s largest assemblage approach this artificial hypothetical climate (**Fig. 4**).

The obvious litmus test for a CA+PF study is whether it can distinguish a random sample of nearest-living relatives from a genuine one. With most Palaeoflora climate tolerance data unreported, we confined ourselves to the >200 nearest-living relatives reported in Quan et al. (2012). Random subsampling from this dataset shows that as the number of nearest-living relatives increases in an assemblage, the reconstructed climate values will eventually converge to a monsoonal (winter precipitation << summer precipitation), warm subtropical climate (*Cwa*) as found today in southern China (i.e. eastern Yunnan, southeastern Sichuan, Guangxi, and Guangdong; **Fig. 6**). This climate is consistently reconstructed by Quan et al. (2012) for assemblages with high numbers of reported nearest-living relatives. The CA+PF and the set of 200 nearest-living relatives used to represent the Eocene of China simply has a high probability to reconstruct a modern *Cwa* climate, even if the assemblage is randomly generated. Furthermore, the threshold for “*statistical significance*”, > 88% ‘coexistence’ (Mosbrugger and Utescher, 1997; Utescher et al., 2014) will be readily reached if more than 20 or 30 NLRs are used, thereby stripping the Coexistence Approach from its only means for detecting erroneous data.

**Figure 6.**
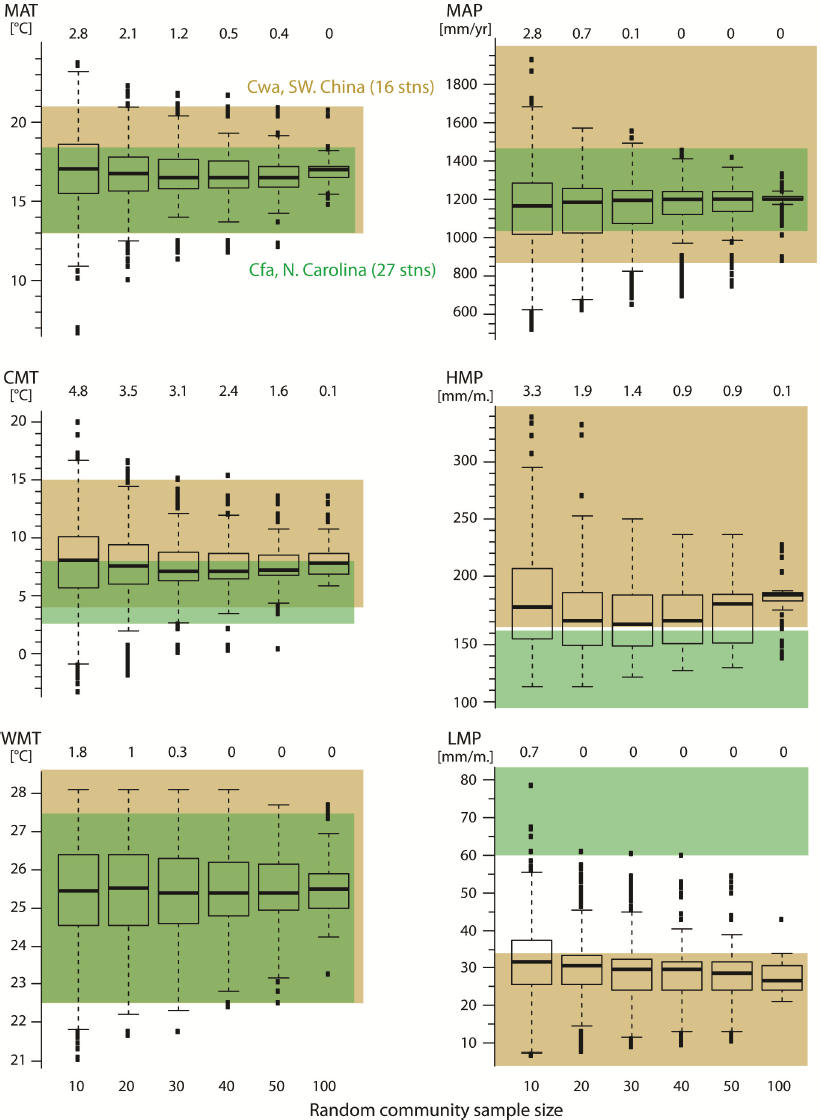
“Yunnan effect” in the data of Quan et al. (2012). Shown are “center values” of coexistence intervals reconstructed for random subsamples of nearest-living relatives (1000 replications per community sample size). Note that with increasing sample size (number of nearest-living relatives), the “center values” of the random assemblages converge to the subtropical *Cwa* climate of modern-day South(west) China and the CA intervals reconstructed for the fossil assemblages (**Figs 1–5, Files S2, S5**). Modern-day East Asian *Cwa* and subtropical *Cfa* North Carolina climates are shown (olive and green backgrounds, respectively). Top numbers in each graph: the percentage of replicates that did not fulfil the >88%-coexistence criterion (cf. Mosbrugger and Utescher, 1997).

## 5 Final assessment of the Palaeoflora database and suggestions for the future of MCR-NLR applications

The underlying data and errors, theoretical and practical, observed in the Quan et al. (2012) study is highly alarming regarding the reliability of CA+PF climate reconstructions as it was conducted by experienced researchers in this field (Utescher et al., 2014). Coexistence intervals are determined by problematic or even invalid nearest-living relatives. The use of the coexistence space of two (unrelated) taxa as estimate for the climate space of a fossil taxon (**Table 5**) that cannot be unambiguously assigned to one of these two taxa violates the fundamentals of the Coexistence Approach (Mosbrugger and Utescher, 1997). The concept of ‘climatic outliers’ and ‘ambiguous intervals’ (Mosbrugger and Utescher, 1997; Utescher et al., 2014), and how they are addressed in CA+PF practise (Quan et al., 2012), are difficult to follow; it neither relates to the background of mutual climate range techniques (Klotz, 1999; Thompson et al., 2012a) nor to the general concept of nearest-living relative approaches. If we assume that all fossils are correctly identified (assumption 1) and associated with a representative nearest-living relative (assumptions 2 and 4) *and* thrived under the same climate than their selected nearest-living relatives (assumption 3), then the latter *must* share a mutual climate range. Thus, 100% ‘coexistence’ is obligatory. Assemblages with < 100% ‘coexistence’ directly indicate that one or several of the basic assumptions of the Coexistence Approach are violated (**Sections 2 and 4.1**).

### 5.1 Obligatory measures regarding Palaeoflora database

As already demonstrated by Grimm and Denk (2012) – who used over 200 modern validation floras with gridded *and* station MAT climate data – the poor quality of, and many errors in, the Palaeoflora database make it unusable for quantitative reconstructions of palaeoclimate using the Coexistence Approach or other techniques. Major drawbacks of the Palaeoflora database are the fixed fossil–nearest-living relative lists, the selection of 4–6 ‘extreme’ climate stations, and logistic problems in curating and updating a restricted-access database.

The fossil–nearest-living relative association lists are a combination of outdated views (e.g. *Rhoipites* → *Rhus*), errors (e.g. Engelhardioidea fossils), and new, possibly more reliable associations. Rather than compiling thousands of taxa at the risk of hundreds, if not thousands, of problematic associations, the Palaeoflora curators should only include carefully checked (using light and scanning electron microscopy for pollen taxa) and reliable associations. To enable reproducibility and correction, each association must be listed with the relevant literature and images of fossils be made available for those that determine coexistence intervals. Ideally, images of all fossil taxa used should be available. This is particularly important for studies using palynological data, where the ‘type status’ may be standardised but not the botanical affinity of these types (compare Stuchlik et al. 2001–2014 with Coexistence Approach literature). It is paramount to provide full taxonomic details of any artificial pollen taxon. Lists like those used provided by Quan et al. (2012) are insufficient. Only through careful documentation can association errors be eliminated. The use of general lists for fossil–nearest-living relative associations is questionable: the application of any MCR-NLR technique should always be accompanied by researchers that have the competence and experience to decide on a meaningful nearest-living relative for a fossil taxon.

The selection of ‘extreme’ climate stations obviously does not fulfil assumption 4 (**Table 2**) of Mosbrugger and Utescher (1997). The problems linked to the use of gridded climate data is no excuse (Utescher et al., 2014) for not incorporating the data generated by Thompson et al. (1999a, b, 2001) in the last 15 years; deviations of up to 5 °C between the WordClim dataset and climate stations do not apply in North America (GWG, pers. obs., 2011–2012). The subjective selection of a climate station inflicts substantially greater errors than the use of gridded climate and distribution data (Grimm and Denk, 2012; this study).

Before studies that rely on the Palaeoflora database can be considered in any way reliable, this database needs to be validated using modern floras as outlined in Grimm and Denk (2012). The users of the Palaeoflora database need to produce hard evidence that the true climate parameters of a modern-day flora fall within the reconstructed climate intervals based on Palaeoflora data. These test reconstructions for modern validation floras must rely on only those taxa that can be traced back in the fossil record. The possible precision and accuracy of any quantitative MCR-NLR approach can only be estimated by applying the same taxonomic resolution problems to the modern-day flora as those observed when investigating palaeo-assemblages (Grimm and Denk, 2012; Thompson et al., 2012a).

### 5.2 Eliminating the main sources of misinformed Coexistence Approach intervals: guidelines for simple and less-biased MCR-NLR techniques

Because of the many unsolved practical (this study) and theoretical (discussed elsewhere, but see e.g. Grimm and Denk, 2012; Denk et al., 2013) issues, we do not recommend the use of Coexistence Approach or other univariate MCR-NLR techniques. Nevertheless, based on our experience with modern and past plant distribution patterns and regional climate anomalies (Denk et al., 2011; Denk et al., 2013; Bouchal et al., 2014; Denk et al., 2014; Grímsson et al., 2014; Velitzelos et al., 2014; Grímsson et al., 2015) and purported Coexistence Approach results (Grimm and Denk, 2012; this study), we can provide guidelines for MCR-NLR methods that will minimise the most imminent biases of CA+PF studies. These guidelines should be followed when using plant fossil assemblages for MCR-NLR techniques (‘capped’ MCR; SC-MCR) are:

1. Avoid family-level NLRs. If generic or subgeneric affinities cannot be unambiguously established, the use of summarised, super-specific/-generic climatic niches may unfortunately be inevitable. But in such cases, the selection of genera (or species) should be meaningful – geographically, systematically and phylogenetically – and reproducible. The application of this can be facilitated if each genus is directly linked to a higher-level taxon, and this information is included in the documentation (see **File S2**). Not providing fixed intervals for subfamilies and families also eliminates the necessity of permanently updating these data when new systematic concepts arise or tolerances are corrected for constituent elements (Utescher et al., 2014). Provided good documentation, the nearest-living relative principle should always be kept in mind when selecting a group of modern taxa as nearest-living relative for a fossil taxon (Denk et al., 2013).
2. Fossils of ambiguous affinity should never be included in taxon-based nearest-living relative approaches. Few, but reliable and meaningful nearest-living relatives *are always better* than many, but unreliable, nearest-living relatives such as used in the study of Quan et al. (2012). **Images for key fossils, the fossils that determine intervals, should be made available**.
3. Avoid using tolerance data of nearest-living relatives that lack proper documentation of distribution data. Referencing all literature that has been used for distribution or bioclimatic data is critical to facilitate error detection. Check if the reconstructed climate is in line with the climate within the present-day distribution area of interval-delimiting nearest-living relatives in order to verify the applicability of the nearest-living relative concept.
4. Rely on values with sensible, not unrealistic precision. In the case of MCR-NLR a good start would be to use categories of 2 °C for temperature and 200 mm/year or 20 mm/month for precipitation parameters. For taxa where exact spatial data available (stand co-ordinates), the error of tolerances can be estimated directly from the data (e.g. by comparison of neighbouring grid cells), which would be particular important for potential nearest-living relatives with a distinct altitudinal distribution.
5. Document all steps, data and decisions to enable current researchers to check your reconstructions and future researchers to confirm the validity of your results as more information becomes available.

### 5.3 Conclusions

This critical appraisal of the first Coexistence Approach study using the Palaeoflora database that provided sufficient documentation to test the reconstructed palaeoclimates demonstrates that this study is severely flawed. This is in contrast to the conclusion of Utescher et al. (2014), who declared that agreement (however anecdotal) with other palaeoclimate reconstruction methods demonstrate that CA+PF produced sound results despite its many sources of errors (“uncertainties”). Using the data provided by Quan et al. (2012), we highlight data errors, methodological errors, and inconsistencies that are undetectable in the last 15 years of CA+PF studies due to a lack of sufficient documentation. Thus, palaeoclimate reconstructions based on the CA+PF method should be discounted until re-analyses and documentation are provided and any future studies should be investigated for the errors highlighted here before trusting the results.

## Background data and supplementary material

An archive including all supporting material (**Files S1** to **S6, Tables S1** to **S4**) is provided for anonymous download at www.palaeogrimm.org/data.

## Acknowledgements

GWG acknowledges funding from the Austrian Science Fund (FWF grant M-1751B16), TD from the Swedish Science Foundation (VR grant 2012-4378), and AJP from the National Research Foundation (RCA13091944022).

## Plate I (following page)

SEM micrographs of all four species of subfamily Taxodioideae (Cupressaceae): *Cryptomeria japonica, Glyptostrobus pensilis, Taxodium mucronatum* and *T. distichium* var. *imbricarium*. **A-D.** Pollen of *Cryptomeria japonica*. **A.** LM, overview. **B.** SEM, overview. **C.** SEM, detail proximal side. **D.** SEM, detail distal side. **E-H.** Pollen of *Glyptostrobus pensilis*. **E.** LM, overview. **F.** SEM, overview. **G.** SEM, detail proximal side. **H.** SEM, detail distal side. **I-L.** Pollen of *Taxodium mucronatum*. **I.** LM, overview. **J.** SEM, overview. **K.** SEM, detail proximal side. **L.** SEM, detail distal side. **M-P.** Pollen of *Taxodium distichium*. **M.** LM, overview. **N.** SEM, overview. **O.** SEM, detail proximal side. **P.** SEM, detail distal side. Scale bars – 10 µm **(A, B, E, F, I, J, M, N)**, 1µm **(C, D, G, H, K, L, O, P)**.

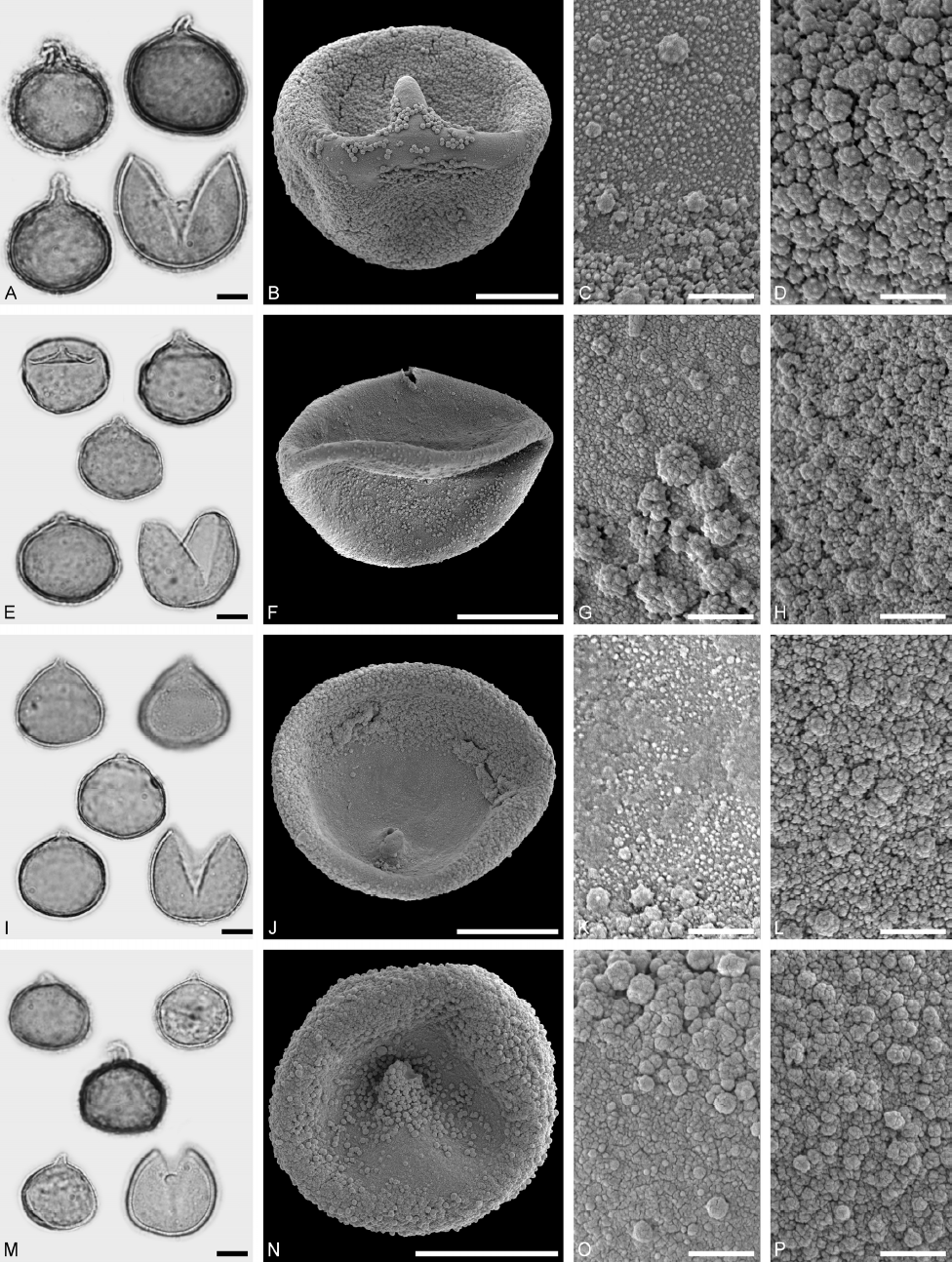

## References

Beug HJ. 2004. Leitfaden der Pollenbestimmung für Mitteleuropa und angrenzende Gebiete. München: Verlag Dr. Friedrich Pfeil.

Böcher J. 1995. Palaeoentomology of the Kap København Formation, a Plio-Pleistocene sequence in Peary Land, North Greenland. Meddelelser om Grønland Geoscience, 33: 1–89.

Bouchal J, Zetter R, Grímsson F, Denk T. 2014. Evolutionary trends and ecological differentiation in early Cenozoic Fagaceae of western North America. American Journal of Botany, 101: 1–18.

Denk T, Grimm G, Stögerer K, Langer M, Hemleben V. 2002. The evolutionary history of *Fagus* in western Eurasia: Evidence from genes, morphology and the fossil record. Plant Systematics and Evolution, 232: 213–236.

Denk T, Grimm GW. 2009a. The biogeographic history of beech trees. Review of Palaeobotany and Palynology, 158: 83–100.

Denk T, Grimm GW. 2009b. Significance of pollen characteristics for infrageneric classification and phylogeny in *Quercus* (Fagaceae). International Journal of Plant Sciences, 170: 926–940.

Denk T, Grimm GW. 2010. The oaks of western Eurasia: traditional classifications and evidence from two nuclear markers. Taxon, 59: 351–366.

Denk T, Grimm GW, Grímsson F, Zetter R. 2013. Evidence from “Köppen signatures” of fossil plant assemblages for effective heat transport of Gulf Stream to subarctic North Atlantic during Miocene cooling. Biogeosciences, 10: 7927–7942.

Denk T, Grimm GW, Hemleben V. 2005. Patterns of molecular and morphological differentiation in *Fagus*: implications for phylogeny. American Journal of Botany, 92: 1006–1016.

Denk T, Grímsson F, Zetter R, Símonarson LA. 2011. Late Cainozoic Floras of Iceland: 15 Million Years of Vegetation and Climate History in the Northern North Atlantic. Heidelberg, New York: Springer.

Denk T, Güner HT, Grimm GW. 2014. From mesic to arid: Leaf epidermal features suggest preadaptation in Miocene dragon trees (*Dracaena*). Review of Palaeobotany and Palynology, 200: 211–228.

Earle CJ. 2010. The Gymnosperm Database.

Eldrett JS, Greenwood DR, Polling M, Brinkhuis H, Sluijs A. 2014. A seasonality trigger for carbon injection at the Paleocene–Eocene Thermal Maximum. Climates of the Past, 10: 759–769.

Elias SA. 2001. Mutual climatic range reconstructions of seasonal temperatures based on Late Pleistocene fossil beetle assemblages in Eastern Beringia. Quaternary Science Reviews, 20: 77–91.

Fang J, Wang Z, Tang Z. 2009. Atlas of Woody Plants in China. Volumes 1 to 3 and index. Beijing: Higher Education Press.

Flora of China. 2014. eFloras: Flora of China.: Missouri Botanical Garden, St. Louis, MO & Harvard University Herbaria, Cambridge, MA.

Flora of North America. 2004 onwards. eFloras: Flora of North America. Missouri Botanical Garden, St. Louis, MO & Harvard University Herbaria, Cambridge, MA.

Gao Z-f, Thomas BA. 1989a. Occurrence of earliest cycads in the Permian of China and its bearing on their evolution. Chinese Science Bulletin, 34: 766–769.

Gao Z-f, Thomas BA. 1989b. A review of fossil cycad megasporophylls, with new evidence of *Crossozamia* Pomel and its associated leaves from the Lower Permian of Taiyuan, China. Review of Palaebotany and Palynology, 60: 205–223.

Greenwood DR, Archibald SB, Mathewes RW, Moss PT. 2005. Fossil biotas from the Okanagan Highlands, southern British Columbia and northeastern Washington State: climates and ecosystems across an Eocene landscape. Canadian Journal of Earth Sciences, 42: 167–185.

Grimm GW, Denk T. 2012. Reliability and resolution of the coexistence approach — A revalidation using modern-day data. Review of Palaeobotany and Palynology, 172: 33–47.

Grimm GW, Denk T, Hemleben V. 2007. Evolutionary history and systematic of *Acer* section *Acer* - a case study of low-level phylogenetics. Plant Systematics and Evolution, 267: 215–253.

Grímsson F, Zetter R, Grimm GW, Krarup Pedersen G, Pedersen AK, Denk T. 2015. Fagaceae pollen from the early Cenozoic of West Greenland: revisiting Engler’s and Chaney’s Arcto-Tertiary hypotheses. Plant Systematics and Evolution, 301: 809–832.

Grímsson F, Zetter R, Halbritter H, Grimm GW. 2014. *Aponogeton* pollen from the Cretaceous and Paleogene of North America and West Greenland: Implications for the origin and palaeobiogeography of the genus. Review of Palaeobotany and Palynology, 200: 161–187.

Hijmans RJ, Cameron SE, Parra JL, Jones PG, Jarvis A. 2005. Very high resolution interpolated climate surfaces for global land areas. International Journal of Climatology, 25: 1965–1978.

Huang T-C. 1972. Pollen flora of Taiwan. Taipeh: National Taiwan University Botany Department Press.

Hutchinson GE. 1957. Concluding Remarks. Cold Spring Harbor Symposia on Quantitative Biology, 22: 415–427.

Jacques FMB, Guo S-X, Su T, Xing Y-W, Huang Y-J, Liu Y-S, Ferguson DK, Zhou Z-K. 2011. Quantitative reconstruction of the Late Miocene monsoon climates of southwest China: a case study of the Lincang flora from Yunnan Province. palaeogeography, Palaeoclimatology, Palaeoecology, 304: 318–327.

Jones GD, Bryant VBJ, Lieux MH, Jones SD, Lingren PD. 1995. Pollen of the Southeastern United States: with emphasis on melissopalynology and entomopalynology. American Association of Stratigraphic Palynologists Foundation - Contributions, Series 30.

Klotz S. 1999. Neue Methoden der Klimarekonstruktion - angewendet auf quartäre Pollensequenzen der französischen Alpen. Tübingen: Institut & Museum für Geologie & Paläontologie [know: Institute for Geosciences], Eberhard Karls University.

Kottek M, Grieser J, Beck C, Rudolf B, Rubel F. 2006. World map of the Köppen-Geiger climate classification updated. Meteorol. Z., 15: 259–263.

Kotthoff U, Greenwood DR, McCarthy FMG, Müller-Navarra K, Prader S, Hesselbo SP. 2014. Late Eocene to middle Miocene (33 to 13 million years ago) vegetation and climate development on the North American Atlantic Coastal Plain (IODP Expedition 313, Site M0027). Climates of the Past, 10: 1523–1539.

Kvaček Z. 2007. Do extant nearest relatives of thermophile European Cenozoic plant elements reliably reflect climatic signal? Palaeogeography, Palaeoclimatology, Palaeoecology, 253: 32–40.

Li T. 2011. Pollen Flora of China Woody Plants by SEM. Bejing: Chinese Cooperation for Promotion of Humanities.

Manchester SR. 1987. The fossil history of the Juglandaceae. St. Louis: Missouri Botanical Garden.

Manos PS, Soltis PS, Soltis DE, Manchester SR, Oh S-H, Bell CD, Dilcher DL, Stone DS. 2007. Phylogeny of extant and fossil Juglandaceae inferred from the integration of molecular and morphological data sets. Systematic Biology, 56: 412–430.

Manum SB. 1962. Studies in the Tertiary flora of Spitsbergen,with notes on Tertiary floras of Ellesmere Island, Greenland, and Iceland. Norsk Polarinstitutt Skrifter, 125: 1–127.

Martin PS, Drew CM. 1969. Scanning electron photographs of southwestern pollen grains. Journal of the Arizona Academy of Science, 5: 147–176.

Miyoshi N, Fujiki T, Kimura H. 2011. Pollen flora of Japan. Sapporo: Hokkaido University Press.

Mosbrugger V, Utescher T. 1997. The coexistence approach – a method for quantitative reconstructions of Tertiary terrestrial palaeoclimate data using plant fossils. Palaeogeography, Palaeoclimatology, Palaeoecology, 134: 61–86.

Müller MJ, Hennings D. 2000. The Global Climate Data Atlas on CD Rom. Flensburg: Universität Flensburg, Institut für Geografie.

Peel MC, Finlayson BL, McMahon TA. 2007. Updated world map of the Köppen-Geiger climate classification. Hydrology and Earth System Sciences, 11: 1633–1644.

Pocknall DT, Crosbie YM. 1988. Pollen morphology of *Beauprea* (Proteaceae): Modern and fossil. Review of Palaeobotany and Palynology, 53: 305–327.

Quan C, Liu Y-SC, Utescher T. 2012. Eocene monsoon prevalence over China: A paleobotanical perspective. Palaegeography, Palaeoclimatology, Palaeoecology, 365-366: 302–311.

Solomon AM, King JE, Martin PS, Thomas J. 1973. Further scanning electron photomicrographs of southwestern pollen grains. Journal of the Arizona Academy of Science, 8: 103–157.

Stevens PF. 2001 onwards. Angiosperm Phylogeny Website. Version 8, June 2007 [and more or less continuously updated since].

Stuchlik L, Ziembińska-Tworzydło M, Kohlman-Adamska A, Grabowska I. B. S, Ważyńska, H., Sadowska A. 2009. Atlas of pollen and spores from the Polish Neogene. Vol. 3. Angiosperms (1). Kraków: W. Szafer Institute of Botany, Polish Academy of Science.

Stuchlik L, Ziembińska-Tworzydło M, Kohlman-Adamska A, Grabowska I. B. S, Worobiec E, Durska E. 2014. Atlas of pollen and spores from the Polish Neogene. Vol. 4. Angiosperms (2). Kraków: W. Szafer Institute of Botany, Polish Academy of Science.

Stuchlik L, Ziembińska-Tworzydło M, Kohlman-Adamska A, Grabowska I, Ważyńska, H., Sadowska A. 2002. Atlas of pollen and spores from the Polish Neogene. Vol. 2. Gymnosperms. Kraków: W. Szafer Institute of Botany, Polish Academy of Science.

Stuchlik L, Ziembińska-Tworzydło M, Kohlman-Adamska A, Grabowska I, Wazyńska H. B. S, Sadowska A. 2001. Atlas of pollen and spores from the Polish Neogene. Vol. 1. Spores. Kraków: W. Szafer Institute of Botany, Polish Academy of Science.

Takahashi M. 1989. Pollen morphology of Celtidaceae and Ulmaceae: reinvestigation. In: Crane PR, Blackmore S, eds. Evolution, Systematics, and Fossil History of the Hamamelidae. Vol. 2. Higher Hamamelidae. Oxford: Claredon Press.

Taylor TN, Taylor EL, Krings M. 2009. Paleobotany – The Biology and Evolution of Fossil Plants. Burlington: Academic Press.

Thompson RS, Anderson KH, Bartlein PJ. 1999a. Atlas of relations between climatic parameters and distribution of important trees and shrubs in North America — Hardwoods. U.S. Geological Survey Professional Paper, 1650–B: 1–423.

Thompson RS, Anderson KH, Bartlein PJ. 1999b. Atlas of relations between climatic parameters and distributions of important trees and shrubs in North America — Introduction and Conifers. U.S. Geological Survey Professional Paper, 1650–A: 1–269.

Thompson RS, Anderson KH, Bartlein PJ. 2001. Atlas of relations between climatic parameters and distributions of important trees and shrubs in North America — Additional conifers, hardwoods, and monocots. u.S. Geological Survey Professional Paper, 1650–C: 1–386.

Thompson RS, Anderson KH, Pelltier RT, Strickland LE, Bartlein PJ, Shafer SL. 2012a. Quantitative estimation of climatic parameters from vegetation data in North America by the mutual climatic range technique. Quaternary Science Reviews, 51: 18–39.

Thompson RS, Anderson KH, Pelltier RT, Strickland LE, Shafer SL, Bartlein PJ. 2012b. Atlas of relations between climatic parameters and distributions of important trees and shrubs in North America—modern data for climatic estimation from vegetation inventories. U.S. Geological Survey Professional Paper, 1650–F: http://pubs.usgs.gov/pp/p1650-f/.

Thompson RS, Anderson KH, Strickland LE, Shafer SL, Pelltier RT, Bartlein PJ. 2006. Atlas of Relations Between Climatic Parameters and Distributions of Important Trees and Shrubs in North America— Alaska Species and Ecoregions. U.S. Geological Survey Professional Paper, 1650–D: 1–342.

Tiffney BH. 2008. Phylogeography, fossils, and northern hemisphere biogeography: The role of physiological uniformitarianism. Annals of the Missouri Botanical Garden, 95: 135–143.

Utescher T, Bruch AA, Erdei B, François I, Ivanov D, Jacques FMB, Kern AK, Liu Y-SC, Mosbrugger V, Spicer RA. 2014. The Coexistence Approach—Theoretical background and practical considerations of using plant fossils for climate quantification. Palaeogeography, Palaeoclimatology, Palaeoecology, 410: 58–73.

Utescher T, Mosbrugger V. 2009. Palaeoflora Database.

Velitzelos D, Bouchal JM, Denk T. 2014. Review of the Cenozoic floras and vegetation of Greece. Review of Palaeobotany and Palynology, 204: 56–117.

Walter H, Lieth H. 1960. Klimadiagramm-Weltatlas, 1. Lieferung. Jena: VEB Gustav Fischer Verlag.

Xu J-X, Ferguson DK, Li C-S, Wang Y-F. 2008. Late Miocene vegetation and climate of the Lühe region in Yunnan, southwestern China. Review of Palaeobotany and Palynology, 148: 36–59.

Zachos JC, Pagani M, Sloan L, Thomas E, Billups K. 2001. Trends, rhythms, and aberrations in global climate 65 Ma to present. Science, 292: 686–693.

Zagwijn WH. 1994. Reconstruction of climate change during the Holocene in western and central Europe based on pollen records of indicator species. Vegetation History and Archaeobotany, 3: 65–88.

